# Intertwined canonical and non-canonical initiation in dual promoters are pervasive and differentially regulate Polymerase II transcription

**DOI:** 10.1101/487496

**Authors:** Chirag Nepal, Yavor Hadzhiev, Estefanía Tarifeño-Saldivia, Ryan Cardenas, Ana-Maria Suzuki, Piero Carninci, Bernard Peers, Boris Lenhard, Jesper B. Andersen, Ferenc Müller

## Abstract

The diversity and complexity of transcription start site (TSS) selection reflects variation of preinitiation complexes, divergent function of promoter-binding proteins and underlies not only transcriptional dynamics but may also impact on post-transcriptional fates of RNAs. The majority of metazoan genes are transcribed by RNA polymerase II from a canonical initiation motif having an YR dinucleotide at their TSSs. In contrast, translation machinery-associated genes carry promoters with polypyrimidine initiator (known as 5’-TOP or TCT) with cytosine replacing the R nucleotide. The functional significance of start site choice in promoter architectures is little understood. To get insight into the developmental regulation of start site selection we profiled 5’ ends of transcripts during zebrafish embryogenesis. We uncovered a novel class of dual-initiation (DI) promoters utilized by thousands of genes. In DI promoters non-canonical YC-initiation representing 5’-TOP/TCT initiators is intertwined with canonical YR-initiation. During maternal to zygotic transition, the two initiation types are divergently used in hundreds of DI promoters, demonstrating that the two initiation systems are distinctly regulated. We show via the example of snoRNA host genes and translation interference experiments that dual-initiation from shared promoters can lead to divergent spatio-temporal expression dynamics generating distinct sets of RNAs with different post-transcriptional fates. Thus utilization of DI promoters in large number of genes suggests two transcription initiation mechanisms targeting these promoters. DI promoters are conserved within human and fruit fly and reflect an evolutionary conserved mechanism for switching transcription initiation to adapt to the changing developmental context. Thus, our findings highlight a novel level of complexity of core promoter regulation in metazoans and broaden the scope for identification and characterization of alternative RNA products generated at shared core promoters.

## Introduction

Transcription is a tightly regulated process initiated by RNA polymerase II (Pol II) in the core promoter region, which is typically −40 to +40 nucleotides with respect to transcription start sites (TSS). There are no universal core promoter elements^1^ as they are diverse in their sequence and functions, and the structure-function relationship of core promoters remains poorly understood. Sequencing of capped RNA 5’ ends by CAGE (cap-analysis of gene expression) revealed that an overwhelming majority of TSSs are anchored by a purine base at the start site (+1 position) and flanked by pyrimidine in the upstream region (−1 position), thus defining consensus Y_-1_R_+1_ (hereafter called YR-initiation) as canonical initiator in mammals^2^ and in teleosts (zebrafish and tetraodon)^3^, suggesting generality of conserved initiator among vertebrates. Analysis of core promoters in *Drosophila melanogaster* (invertebrates) revealed a related but more motif-like TC_-1_A_+1_GT initiator sequence^4,5^. In contrast, transcription initiation of translation-associated genes (ribosomal proteins, snoRNA host genes, translation initiation and elongation factors) is anchored by C_+1_ (cytosine) and flanked by a polypyrimidine stretch^6-11^ (hereafter called YC-initiation). These non-canonical initiators have previously been termed 5’-TOP (terminal oligo-polypyrimidine) in mammalian systems or TCT initiators in *Drosophila*^12^ and these YC initiation-dependent genes were shown to be conserved in zebrafish^3^. *Drosophila* ribosomal protein genes with TCT promoters are recognized by a TFIID-independent transcription initiation mechanism and mediated by the TATA-binding protein (TBP) family member TBP-related factor 2 (TRF2), but not TBP^13^. These results suggest that the non-canonical initiation is specialized for a subset of genes and facilitates a non-canonical initiation complex formation with distinct proteins from that of TBP and TFIID and likely reflecting distinct regulation of transcription initiation^14^. However, it is unknown, why such a non-canonical initiation has evolved and has been maintained in evolutionary distant species. Important insight into potential functional significance of the non-canonical initiation is emerging from studies investigating target genes of mTOR pathways that are translationally regulated^15,16^, and are enriched in 5’-TOP/TCT initiator. The 5’-TOP initiator is defined by a minimum of 4-15 pyrimidine sequences^17^. The polypyrimidine stretch proximal to the 5’ end of these genes is a target for translation regulation and has been suggested to serve as a target mechanism for oxidative and metabolic stress or cancer-induced differential translational regulation by the mTOR pathway^15,16,18-20^. The existence of 5’-TOP/TCT promoters raises the questions of how widespread non-canonical initiation is and what is its relationship with canonical initiation.

We have generated CAGE datasets^3^ in zebrafish and profiled all transcription initiators during embryogenesis from the maternal to zygotic transition (MZT) and then through organogenesis. We have extended the detection of YC-initiation in zebrafish to thousands of genes, and made constellation observation of pervasive co-occurrence of YR-initiation and YC-initiation events in shared core promoter. We performed a comprehensive and unbiased analysis of TSSs in promoters and characterized the features and roles of non-canonical initiation by a systematic survey of the base composition within the TSSs in CAGE datasets^3^. This analysis led us to uncover non-canonical YC-initiation in thousands of genes that are proximal to or intertwined with the canonical YR-initiation in the same core promoter region, thus revealing thousands of what we term dual-initiation (DI) promoter genes. We provide multiple lines of evidence for the functional relevance of dual-initiation, such as sequence composition, differential usage of initiators during development, differential response of initiators during translation inhibition and selective association of snoRNA biogenesis, which is predicted to be processed by splicing from introns of the YC-initiation products of dual-initiation genes. We thus demonstrate that the two initiation types within dual promoters represent composite of promoter architectures and reflect on two regulatory functions, generating distinct sets of RNAs with different post-transcriptional fates. Our findings highlight a novel level of complexity of core promoter regulation during development and broaden the scope for functional dissection of overlaid promoter architectures that act in the complexity of the developing embryo.

## Results

### Non-canonical YC-initiations are pervasively intertwined with canonical YR-initiations

To comprehensively map non-canonical initiation events at single nucleotide resolution, we analyzed the start base distribution of (m)RNA 5’ ends by pooling CAGE Transcription Start Sites (CTSSs) with at least 1 tag per million (TPM) detectable across 12 stages during zebrafish embryo development ^3^(**Figure 1a**). Majority of CTSSs (71.6%) have canonical (Y_-1_R_+1_) start sites (**Figure 1a**; **Supplementary Figure 1a**). The remaining CTSSs have been excluded from further analysis as they include RNA start sites with a well-characterized GG dinucleotide associated with post-transcriptional processing products independent from transcription initiation^3^ and therefore do not reflect true transcription start sites. Furthermore, we have excluded CAGE signals which represent Drosha-processing sites on pre-miRNAs^21^ and snoRNA 5’-end capping events^22^. Importantly, a substantial proportion of TSSs possess the non-canonical pyrimidine initiation (labeled Y_-1_C_+1_ in **Figure 1a**). Majority of YR-initiation (85.97%) and YC-initiation (83.05%) sites mapped within the expected promoter region of ENSEMBL transcripts (500 bases upstream and 300 bases downstream) and thus, support detection of true transcription initiation products. YR-initiation and YC-initiation are highly reproducible across replicates (**Supplementary Figure 1b**). For downstream analysis, we retained only those robustly detected transcripts that are transcribed in at least 2 developmental stages and whose promoter expression level is at least 3 TPM. At this filtering threshold, 4201 promoters have YC-initiation and 12056 promoters have YR-initiation (**Supplementary Table 1**). Intersection analysis of gene promoters revealed that 50 (1.19%) genes carry only YC-initiation and 7905 (65.5%) genes have only YR-initiation, thus regulated by a single type of initiator. However, the majority of YC-initiation site-containing promoters (98.81%) also carry YR-initiation sites (**Figure 1a**; Venn diagram). This novel class of promoters have collectively called dual-initiation (DI) promoters (**Figure 1b**). The DI promoters identified by CAGE were also confirmed by independent analysis of capped mRNA sequencing at prim 5 stage of development (24h post fertilization), which, though less sensitive than CAGE, has demonstrated hundreds of cases of dual-initiation events and demonstrated statistically significant overlap with CAGE detected dual-initiation promoter genes (**Supplementary Figure 1c**).

**Figure 1.**
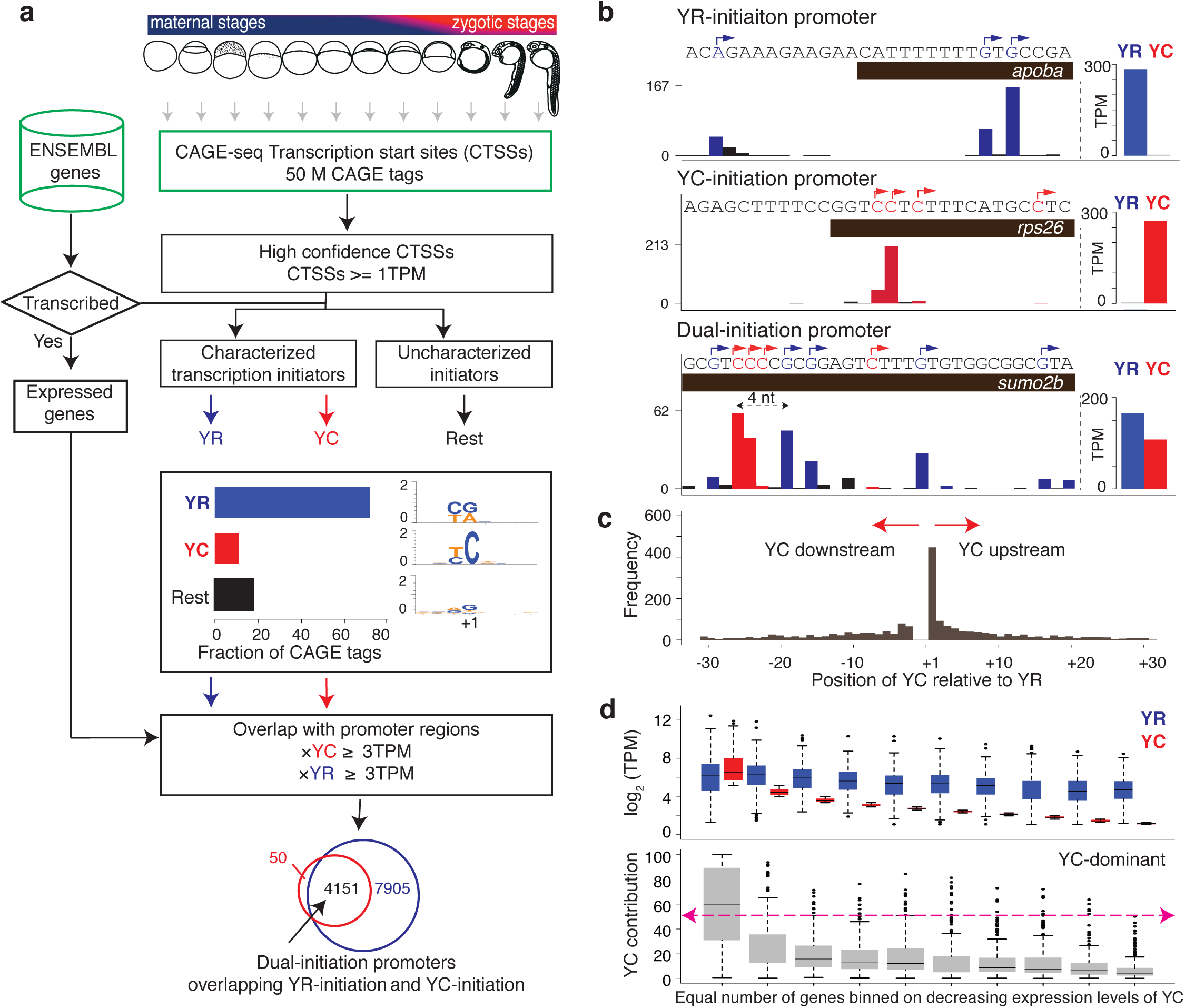
Intertwined canonical initiator (YR) and non-canonical initiator YC (alias as TCT/5’TOP) within the same core promoter. **(a)** A systematic pipeline for identification of canonical (YR) and non-canonical (YC) initiators in the zebrafish developmental promoterome. CTSSs are classified into known YR and YC initiators based on CAGE transcription start sites (CTSSs). **(b)** UCSC browser views with CAGE data from prim 5 stage to illustrate examples of YR-initiation (apoba), YC-initiation (rps26) promoters along with a gene promoter with intertwined YR-initiaions and YC-initiations (sumo2b). YR-initiations and YC-initiations are shown in blue and red colors respectively. Barplot on the right shows the sum of expression levels of YR-initiations and YC-initiations. Highest CTSS represents the dominant transcription start site. The distance between dominant YR and YC in sumo2b is four nucleotides. (**c**) Position of dominant YC-initiation relative to dominant YR-initiation. (**d**) Contribution of YC-initiation with respect to YR-initiation expression levels in prim 5 stage. The 4151 genes with dual-initiation are sorted according to YC expression levels and grouped into 10 % bins. Abbreviations: TPM, tags per million.

For all dual-initiation promoter genes, we summed the expression levels of all YR and YC components and genes were classified as either YR-dominant or YC-dominant depending upon the TPM levels of their YR and YC components. The exemplified *sumo2b* gene (**Figure 1b**) has a higher total level of YR-initiations than YC-initiation, thus classified as a YR-dominant gene. We then used the highest expression level of YR and YC CTSSs and determined the position of dominantly used YR and YC TSS. The YR-dominant TSS is located 4 nucleotides downstream to the YC-dominant TSS in the exemplified *sumo2b* gene (**Figure 1b**). The distance between dominant YR-initiation and YC-initiation of all DI promoters at prim 5 stage fall mostly within 30 bases and there is some degree of preference for YC 1 nt upstream of YR (**Figure 1c**). This close proximity between the two types of initiations suggest that the initiation machinery or machineries involved in controlling transcription of these transcripts recognize the same core promoter region. Comparing the expression levels of YR and YC components revealed that the contribution of YC-initiation to the total activity of dual-initiation promoters tends to be relatively small (**Figure 1d; Supplementary Figure 1d**), resulting only in a small portion (8.25%; n=251) of genes as YC-dominant in prim 5 stage (**Figure 1d**). This observation may explain why the non-canonical YC-initiation events largely have been missed in previous studies, which focused on the single dominant TSSs. However, YC-initiation can be dominant over YR-initiation in individual genes even at lowly expressed promoters (**Figure 1d; Supplementary Figure 1d)**. In conclusion, we show that non-canonical YC-initiation events are pervasively intertwined with canonical YR-initiation and occur within a small physical distance within the same core promoter regions.

### Features of dual-initiation gene promoters

Translational-associated genes such as ribosomal proteins, translation initiation/elongation factors and small nucleolar RNA (snoRNA) host genes are transcribed by 5’-TOP/TCT initiators, thus we asked whether their zebrafish homologs possess single or dual-initiation. The annotation of zebrafish snoRNAs is not comprehensive, therefore we analyzed a size selected RNA library^23^ enriched for full-length snoRNA length (18-250 nt) and annotated 176 novel zebrafish snoRNAs (**Supplementary Table 2**). Intersection of the expressed genes from the above listed gene-families revealed that most of these genes carry dual-initiation sites (**Figure 2a**). Gene ontology (GO) analysis of DI promoter genes revealed an enrichment of translation machinery components (translation, translation elongation and translation termination), co-translational proteins targeting to membrane, RNA stability and nonsense mediated decay (**Figure 2b; Supplementary Table 3**). Enrichment of ribosome-related functions is consistent with previous studies describing YC-initiation^17,24^, associated with such genes while our findings reveal a novel, dual-initiation featuring these promoters (**Figure 2a**). Excluding translation-associated genes from the query list revealed an enrichment of additional unexpected GO terms such as mRNA splicing via spliceosome, telomerase RNA localization, chromosome organization and mitotic cell cycle (**Figure 2b**; **Supplementary Table 3**). In contrast, YR-only initiator genes are enriched for GO terms related to morphogenesis, pattern specification and embryonic development (**Figure 2b**) characteristic of the prim 5 stage of development and highlight the functional distinction of core promoter architectures.

**Figure 2.**
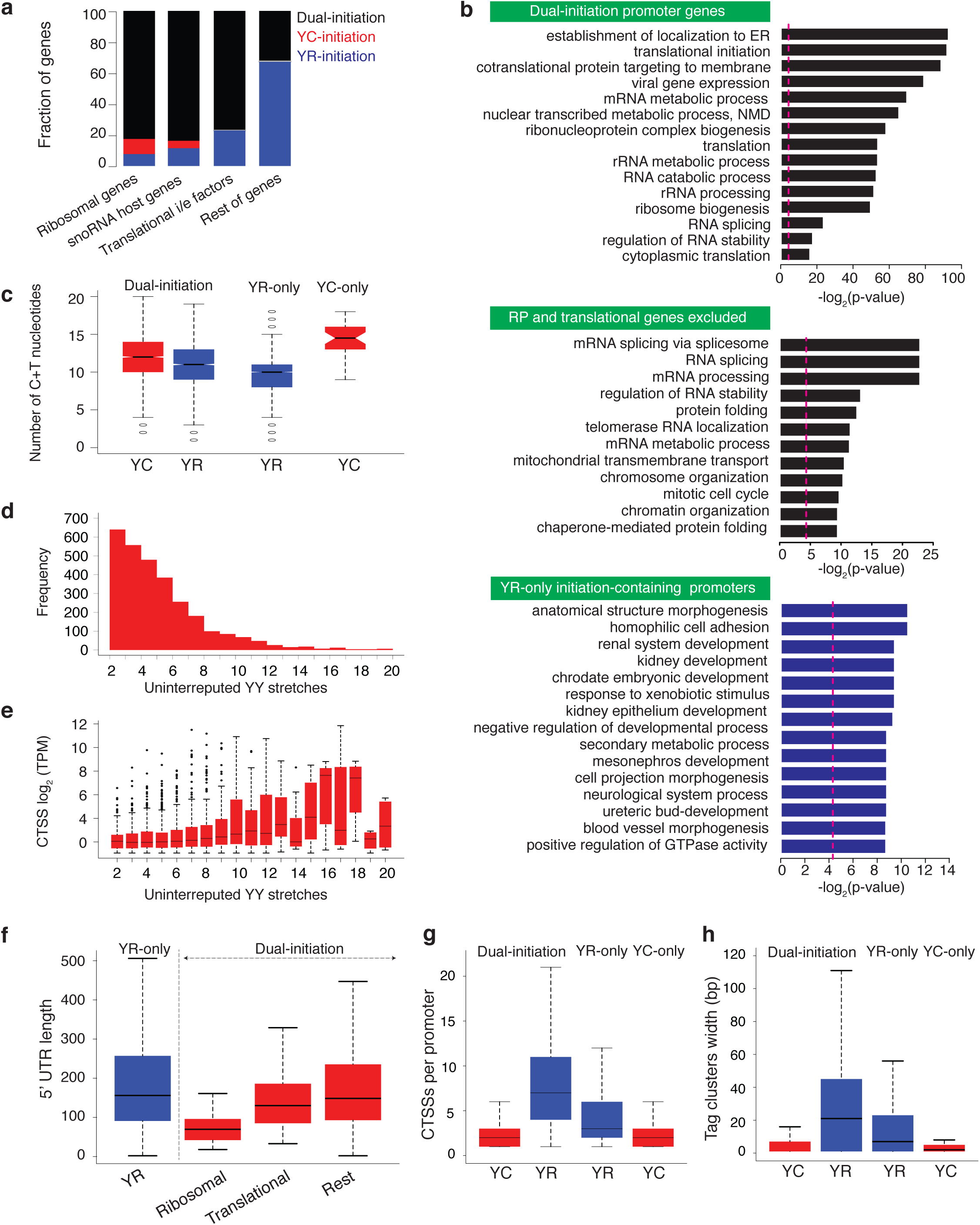
Characteristic features of dual-initiation and single initiation promoter genes. **(a)** Intersection of translation-associated gene families as indicated with single/dual-initiation promoter genes. (**b**) Gene ontology (GO) categories of single and dual-initiation promoter genes clustered as indicated in green fields. (**c**) Sequence composition around dominant YR-initiation and YC-initiation sites of single/dual-initiation promoters. (**d,e**) Presence of polypyrimidine stretches in DI pormoters. X-axis indicates the length of uninterrupted pyrimidine stretch with respect to YC-initiation frequency (**d**) and expression levels of YC-initiation sorted by increasing frequency of uninterrupted polypyrimidine stretches (**e**). (**f**) 5’ UTR length of dual-initiation and single initiation YR genes. (**g**) Frequency of CTSS in single/dual-initiation promoter genes (**h**) Tag cluster width of single/dual-initiation promoter genes.

Sequence composition around (10 nucleotides) dominant TSSs of both initiation sites revealed higher fraction of pyrimidine sequence adjacent to the YC-initiation (**Figure 2c**), predominantly with an uninterrupted stretch of at least 4 pyrimidines (**Figure 2d**), a characteristic feature of the 5’-TOP motif (reviewed in^17^). We find that the longer an uninterrupted pyrimidine stretch around YC-initiation, the higher is the expression level of dominant YC CTSSs (**Figure 2e**). Translation-associated genes have a longer stretch of pyrimidine sequence (**Supplementary Figure 2a**), which is in agreement with the stringent definition of translationally regulated 5’-TOP mRNAs^15^. Dual-initiation promoter genes have shorter 5’-UTR length as compared to single initiation YR promoters (**Figure 2f**), which may reflect efficient translation as transcripts with longer 5’UTR tend to have lower translational efficiency^25^.

Next, we sought to define the promoter features of YR-components and YC-components of dual-initiation promoters. CAGE defined TSSs have revealed 3 main classes of promoter shapes, namely broad peak, sharp peak and bimodal peaks^2^, and 5’-TOP/TCT promoters were primarily associated with sharp peak promoters of highly expressed genes^1^. To explore features of promoter shapes of dual-initiation genes, we first calculated the number of CTSSs and observed that dual-initiation genes have higher number of YR-initiation sites (an average of 6 CTSSs) as compared to their YC constituent (an average of 2 CTSSs) or compared to the YR-only genes (an average of 3 CTSSs) (**Figure 2g**). Accordingly, YR component of dual-initiation promoters is typically defined by a broad peak, while YC-initiation events appear mostly sharp (**Figure 2h**). We then asked if positionally constrained motifs characteristic of known promoter architectures can be assigned to either YC and YR-initiation events in DI promoters. We have plotted YR, YY, SS, WW (Y=C/T; R=A/G; S=C/G; W=A/T) dinucleotides and positionally constrained motifs (TATA box, GC box and CCAT motif) with respect to YR and YC-initiation events at fertilized egg and at prim 5 stage. The WW dinucleotide (W-box motif) present in most promoters in zebrafish^26^ is enriched in both initiators in the fertilized egg but depleted in prim 5 stage (**Supplementary Figure 2b,c**). The finding that YC-initiation is associated with positionally constrained motif previously described for YR-initiation supports YC-initiation detection as indicator of promoter function. Moreover, we have detected similar developmental utilization of sequence determinants of YC transcription start site choice to that previously described for YR-initiation^26^. However, TATA box, CCAT motif and GC box were not enriched with either initiation events in both stages (**Supplementary Figure 2b-c**). Thus, we conclude that YR-initiations peaks of dual-initiation genes are generally broad, while YC-initiations are sharp, however these differences are not reflected in observable differences in the frequency of positionally constrained motifs. Taken together, our results collectively demonstrate the pervasive nature of YC-initiation in the genome which is characteristic not only to translation-associated genes but to previously unappreciated GO categories and often feature TOP promoter-like pyrimidine stretches. These observations suggest that the DI promoter is a novel promoter classification category widely used in the zebrafish genome and which appears to be a composite of canonical and 5’-TOP/TCT promoter features.

### Differential regulation of YC-initiations and YR-initiations in DI promoters during embryogenesis

We have previously shown that two distinct and independently regulated promoter sequence codes such as the W-box and +1 nucleosome positioning signals are often overlaid in individual promoters and used differentially during the maternal to zygotic transition of embryo development^26^. The existence of such overlapping sequence codes, together with the observation that TCT promoters and canonical initiator may be regulated by different initiation complexes^12,13^ prompted us to hypothesize that intertwined YR-initiation and YC-initiation events may represent differential regulatory principles. Thus divergent regulatory inputs may target dual-initiation promoters, and lead to divergent transcriptional regulation during embryo development. Therefore, we asked about the relationship between the expression dynamics of YR-initiation and YC-initiation during early embryo development. We performed self-organizing map (SOM) clustering between YR and YC expression levels for each gene, and observed the typical zebrafish developmental expression profiles, characterized by two opposing trends. A typical maternal dominant trend includes mRNA expression at early stages originating from the oocyte, which is removed by RNA degradation after zygotic genome activation manifesting as loss of expression typically after 6^th^ to 9^th^ stages (**Supplementary Figure 3a,** e.g. panels of first column). An opposite zygotic dominant trend features low or no maternal activity followed by the zygotic activation, which also appears as an increase in expression after the 6^th^ to 9^th^ stages of 12 stages analyzed. Additional trends variations in maternal to zygotic activity of YC and YR have also been detected (**Supplementary Figure 3a**). Most clusters show similar expression dynamics, while differences may have been masked by the pooling of many genes. Nevertheless, several clusters are characterized by distinct profiles for YR and YC components (**Figure 3a),** where the YR component is expressed both maternally and zygotically, whereas the YC component is either zygotic (top row) or maternal only (bottom row). Correlating the expression levels between YR-initiation and YC-initiation during embryogenesis revealed that a majority of genes (71.4%; n=2947) are positively correlated (r >= 0.5), while a small but distinct proportion (7.5%; n=312) of genes show YC and YR components negatively correlated (r <= −0.5) (**Figure 3b**; **Supplementary Table 4**). To understand the origin of negative correlation in regulation, we plotted the expression profiles of these 312 genes and observed two groups with divergent regulation of YR-initiation and YC-initiation during MZT (**Figure 3c**). YR and YC components show opposite maternal zygotic dominance indicating they are distinctly subjected to maternal mRNA degradation and corresponding zygotic transcription activation^26-28^ (**Figure 3d**). Genes in the top cluster predominantly use YR-initiation during maternal stages, in contrast YC-initiation gets dramatically upregulated at the zygotic genome activation after the mid blastula transition (**Figure 3c,d**). This trend is demonstrated by translation elongation factor (*eef1g*) gene promoter (**Figure 3e**), the human homolog of which is transcribed by a non-canonical YC-type initiator^17^. The other negatively co-regulated cluster (bottom cluster in **Figure 3c)** is primarily driven by YC-initiation in maternal stages and by increased YR-initiation in zygotic stages (**Figure 3d**), as exemplified by the initiation profile of the *psmd6* gene (**Figure 3f)**. These results indicate that YR-initiation and YC-initiation are widely used in development and not specific to maternal or zygotic stages. However, they are selectively used for individual genes, which suggests that these genes can respond to differential regulatory inputs. Taken together, the expression dynamics within these 312 dual-initiation promoters indicate independent regulation of YR-initiation and YC-initiation components, which is markedly apparent during the dramatic overhaul of the transcriptome at the MZT.

**Figure 3.**
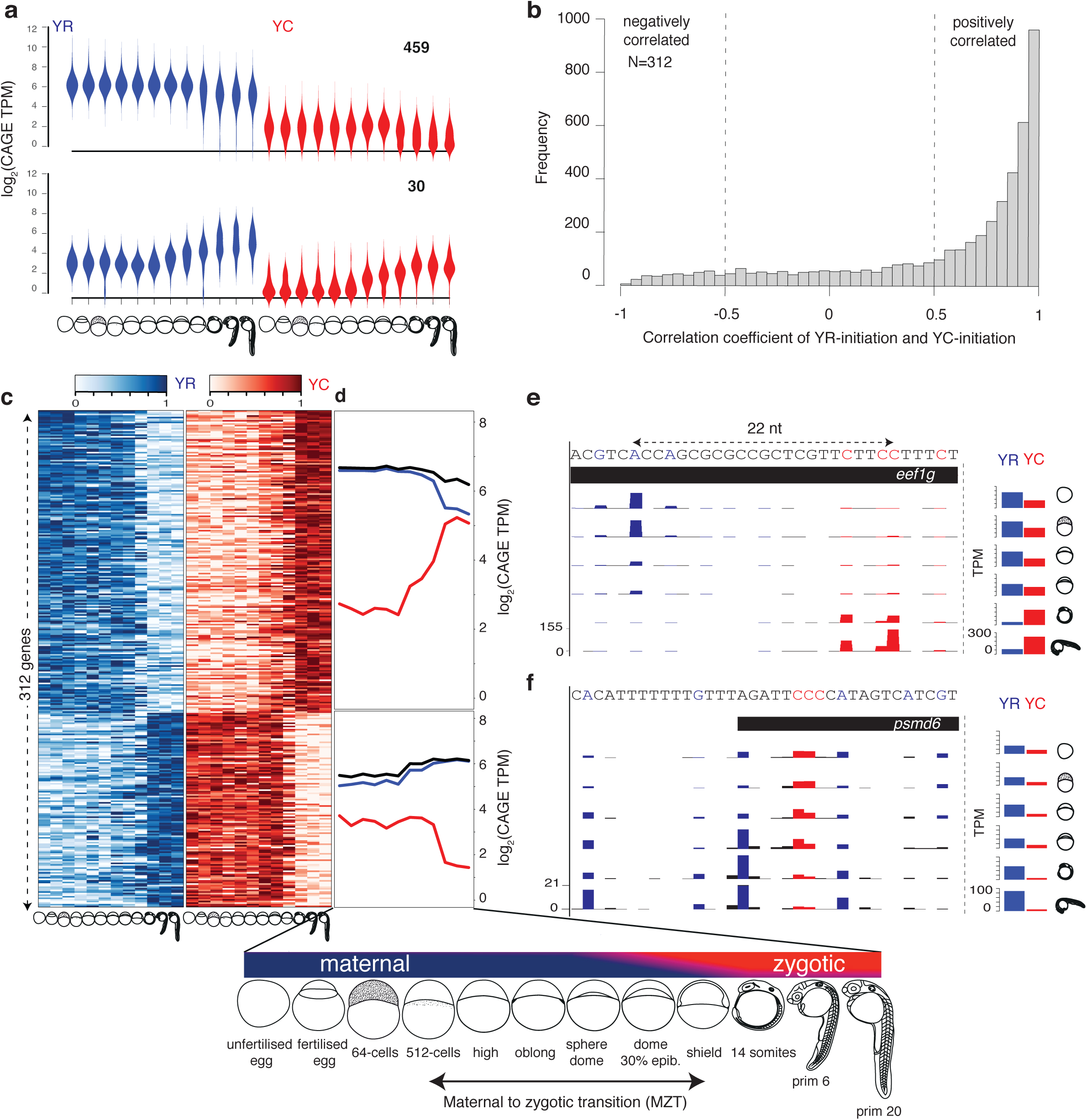
Maternal to zygotic transition of YR-initiation and YC-initiation demonstrates selective promoter utilization in early development. **(a)** Violin plot of expression profiles (tags per million) of YR and YC components of genes during embryo development. X-axis represents developmental stages as indicated. Y-axis indicates the expression levels. Blue and red colors indicate YR and YC components respectively. Numbers indicate genes in the cluster. (**b**) Correlation of expression levels of YR-initiation and YC-initiation during maternal and zygotic stages. X-axis indicates genes binned according to their correlation coefficient. Genes with correlation coefficient (r >= 0.5) are positively correlated and genes with correlation coefficient (r <= −0.5) are negatively correlated. (**c**) Heatmaps show the gene expression profiles of YR-initiations and YC-initiations of 381 negatively correlated genes. Expression values are scaled (row wise) between 0 to 1, separately for YR and YC. Genes are ordered into two groups based on shift from YR to YC (top) and YC to YR (bottom) during maternal and zygotic stages and sorted based on decreasing order of negative correlation in each group. (**d**) Averaged expression level of YR-initiation and YC-initiation across clustered group of genes. (**e,f**) UCSC genome browser views of CTSSs for the eef1g and psmd6 gene promoters. YR-initiation and YC-initiation events are shown in blue and red colors respectively. Barplots on the right shows the sum of CTSSs of YR-initiation and YC-initiation events respectively.

### YC component of dual-initiation promoter genes regulates snoRNA expression

snoRNAs are transcribed by host gene promoters and are spliced out from introns of primary transcripts and subsequently form a riboprotein complex^29^. Thus snoRNA host genes may carry two functional entities, snoRNA genes and their coding or non-coding host gene. Interestingly, a non-coding host gene (*GAS5*) of snoRNA^6^ has recently shown to have an additional function in maintaining nodal signaling^30^. In contrast to previous studies in mammals that described snoRNA host genes being transcribed by YC-initiation (5’-TOP/TCT), we showed that zebrafish snoRNA host genes are characterized by dual-initiation (**Figure 2a**). These observations raise the question, whether the dual function of snoRNA host genes is decoupled by YR-initiation and YC-initiation and whether the two initiation events contribute selectively to distinct RNA fates. Indeed, it was previously shown that a 5’-TOP promoter element determines the specific ratio of snoRNA to mRNA production and an artificial canonical YR-initiation containing Pol II promoter is incompatible with the efficient release of snoRNA^11^. The dramatic transition of maternal and zygotic transcriptomes and the uncovered differential regulation of YC-initiation and YR-initiation at MZT provides an opportunity to address whether YR and YC components of snoRNA host genes are differentially regulated. We thus hypothesized that potentially divergent expression dynamics of YR and YC derived transcripts during MZT could be informative to separate 5’ end of the source RNA for embedded snoRNA genes in dual-initiation promoter host genes. To this end, we plotted the expression levels of both YR and YC components of 97 snoRNA host genes (containing 249 snoRNAs) and the expression of snoRNAs^23^ at the corresponding developmental stages (**Supplementary Figure 4a**). The majority of snoRNA host genes are maternally deposited, and both YR and YC activity as well as snoRNA expression are generally increased after activation of zygotic transcription (**Supplementary Figure 4a**). Correlation of expression levels of snoRNAs with YR and YC components revealed stronger correlation of the YC component with the temporal dynamics of snoRNAs (**Figure 4a**), suggesting YC-initiation to be the likely source for snoRNA host RNA species.

**Figure 4.**
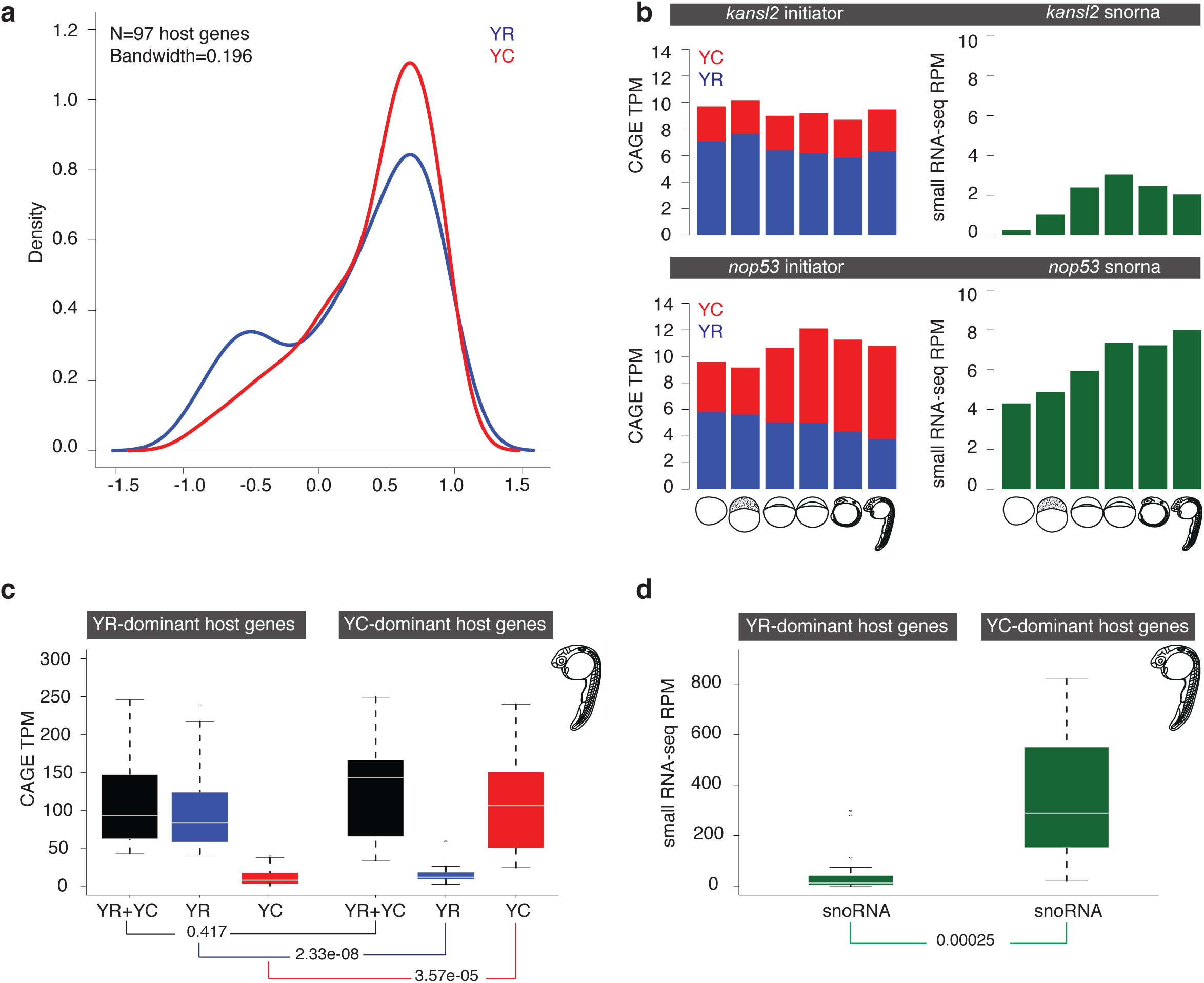
Correlation of expression levels of YR and YC components of snoRNA host genes with that of snoRNA expression levels. **(a)** Correlation of expression levels of YR-initiation and YC-initiation events with snoRNA expression levels across six developmental stages. (**b**) Stacked bar plot of TPM expression levels of YR (blue) and YC (red) components of kansl2 and nop53 genes obtained by CAGE. The expression levels of snoRNA (dark green) calculated from small RNA-seq data are represented in reads per million. Developmental stages are indicated at the bottom. (**c**) Box plot of TPM expression levels of YR-initiation (blue) and YC-initiation (red), along with combined (black) expression levels of YR-initiation and YC-initiation during prim 5 stage. Based on the dominant expression levels of YR-initiation and YC-initiation, host genes are classified as YR-dominant or YC-dominant genes. (**d**) Box plot of expression levels of corresponding snoRNAs (green) from YR-dominant and YC-dominant host genes.

To further investigate the observed correlation between snoRNA expression with YC-initiation, we selected two host genes (*kansl2* and *nop53*) whose overall expression levels are comparable but have varying levels of YR and YC components. The snoRNA host gene *kansl2* has a dominant YR-initiation and a minor YC-initiation, while its snoRNA expression levels is low throughout development (**Figure 4b**). On the other hand, the *nop53* host gene predominantly shows the usage of YC-initiation in zygotic stages and corresponding similar dynamics of snoRNA expression levels (**Figure 4b**). We then analyzed snoRNAs expression levels in relation to the expression of YR and YC components of their host genes at the prim 5 stage, by which time post-transcriptional effects of maternal mRNA clearance are eliminated. We classified 97 snoRNA host genes into YR-dominant (N=44) and YC-dominant (N=53) groups and plotted the expression levels of YR-components and YC-components of host promoter and the corresponding snoRNAs (**Supplementary Figure 4b**). The expression levels of snoRNAs from YC-dominant genes is significantly higher (t test; p=0.00037). Since the overall expression levels of YC-dominant genes is significantly higher than YR-dominant genes (**Supplementary Figure 4b**), higher expression levels of snoRNA is expected, and thus it is difficult to distinguish the contribution of two initiators. Thus, we sought to analyze snoRNA expression levels only in those host genes whose overall expression levels are comparable but have significantly varying contribution of YR and YC components between YR-dominant (N=24) and YC-dominant (N=16) genes (**Figure 4c**). Though overall expression levels are comparable, snoRNAs expression levels are significantly higher (t test; p=0.00025) in YC-dominant genes (**Figure 4d**). Taken together, we provide evidence for divergent developmental regulation of two intertwined initiators in snoRNA host genes. Furthermore, the correlation analysis of temporal and expression levels suggests that the YC-initiation better explains snoRNA expression than the YR-initiation. Nevertheless, the localization of snoRNAs in many ribosomal and translation factors suggests that snoRNAs are produced together with the translation and rRNA biogenesis protein machinery encoded by their host genes and hence they are likely also co-regulated.

### Differential expression and localization of snoRNA and host RNA in zebrafish embryos

The above results suggest that snoRNA host RNAs may be divergently expressed. However, their temporal expression dynamics may not reveal the full extent of differential RNA regulation which emerge from dual-initiation promoter genes. Therefore, we investigated the spatial expression patterns of two newly annotated snoRNAs (**Supplementary Table 2**) embedded in the intron of host gene *nanog* (**Figure 5a**) and *dyskerin* (*dkc1*) respectively (**Figure 5b**). The snoRNA in *nanog* is conserved among teleosts (**Figure 5a**) and is validated by RT-PCR (**Supplementary Figure 5a**). The host gene *nanog* is a transcription factor that regulates genome activation during early zebrafish development^28,31^ with no reported function in rRNA biogenesis. The *nanog* gene carries YR-dominant initiation and low level of YC-initiation (**Supplementary Figure 5b**), with corresponding low level of snoRNA expression. An antisense probe raised against the snoRNA was detected in some but not all nuclei of zebrafish embryos at the sphere stage, whereas an exonic probe detects *nanog* distinctly in the cytoplasm in most cells, indicating the differential transcriptional and/or post-transcriptional fates of the two RNA products generated by the dual-initiation promoter (**Figure 5c-f**).

**Figure 5.**
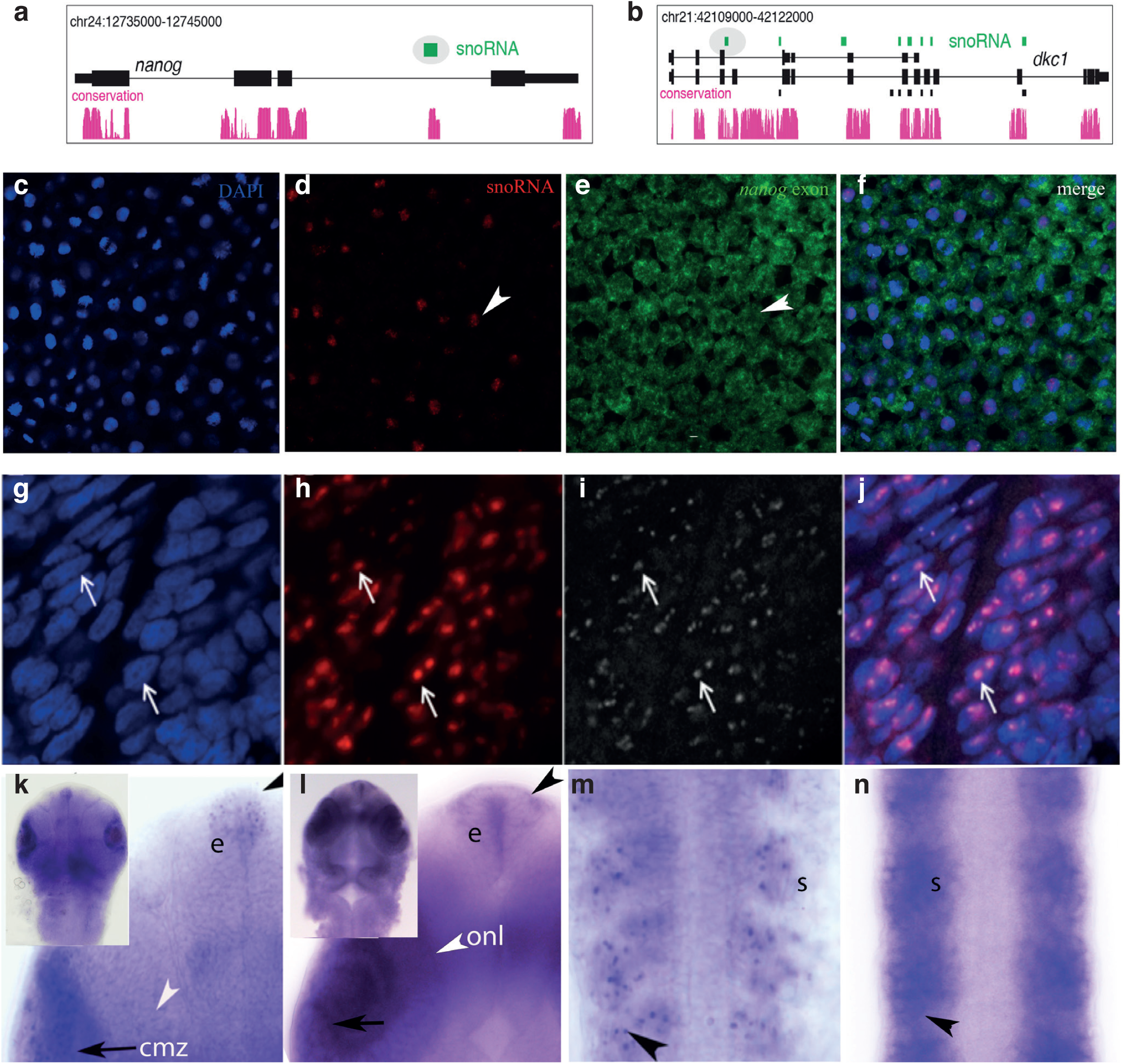
Localization of snoRNAs and host mRNA products in the embryo. (**a-b**) A UCSC browser showing annotated snoRNAs (green) in the introns of nanog and dyskerin (dkc1). Ensembl annotated genes and snoRNAs are shown as black tracks. Teleost sequence conservation tracks are shown in magenta. Two snoRNAs selected for expression analysis are highlighted in oval. (**c-e**) in situ hybridization in whole mount zebrafish embryos at the 30% epiboly stage with probes detecting nanog coding exon and the snoRNA gene embedded in nanog. Probes detected are marked in the panels. **g-j**) In situ hybridization with snoRNA probe from the dyskerin gene is detected in the nucleoli of somites (**g**, overlay in **j**) as indicated by simultaneous immunohisto-chemical detection of fibrillarin (**h**, overlay in **m**). (**k**) snoRNA gene probe detecting snoRNA expression in the ciliary marginal zone of retina (cmz, arrow), epiphysis (e, black arrowhead) and somites (s, arrowhead). (**l-n**) Exon probe of dkc1 indicate cytoplasmic expression in ciliary marginal zone (cmz in **l**) across the retina including the outer nuclear layer (onl, white arrowhead in **k**), epiphysis (e, black arrowhead in l) and somites (s, arrowhead in **n**). Inserts in **k** and **l** show head from dorsal view from which magnified view is cropped.

A snoRNA is produced in several copies from introns of the *dyskerin* (*dkc1*) gene and expected to have shared expression pattern with its host gene given their shared role in pseudouridinylation of ribosomal RNA. We validated one of the novel snoRNAs by RT-PCR (highlighted in oval shape in **Supplementary Figure 5c**). The *dkc1* gene carries YR-dominant initiation in both maternal and zygotic stages (**Supplementary Figure 5d**), while 3 of 4 minor YC-initiation sites become activated higher in zygotic stages (**Supplementary Figure 5c,e**). Expression of the snoRNA by in situ hybridization in whole mount embryos revealed co-localization with Fibrillarin in highly expressing tissues thus, verifying the expected nucleolar expression profile (**Figure 5g-i**). Furthermore, selective expression of snoRNA in nucleoli were detected as speckles in nuclei of a subset of cells at prim 5 stage notably in the epiphysis, somatic muscle cells, and the ciliary marginal zone of the eye. The host RNA *dkc1* exonic probe was detected ubiquitously in the cytoplasm with elevated activity in overlapping (e.g. epiphysis, ciliary marginal zone of retina) as well as non-overlapping domains (e.g. outer nuclear layer of retina) with snoRNA probe (**Figure 5j-m**). Taken together, these two examples suggest that besides the expected differential subcellular localization of host gene products and embedded snoRNAs they are also activated in partially overlapping domains of the embryo, which is consistent with potential divergence in transcriptional regulation of products from the same core promoter.

### Differential fates of YR-initiation and YC-initiation products during translation inhibition

SnoRNA host genes are selectively subjected to nonsense mediated decay (NMD), shown by blocking NMD with translation inhibitor cycloheximide, which led to stabilization of several (*UHG* and *GAS5*)^6,32^, but not all (e.g. *U17HG*^7^, *U87HG*^33^, *rpS16*^6^) snoRNA host genes. This result suggests differential stabilization of host RNAs due to differential association of snoRNA host mRNAs with translating ribosomes^7^. We asked whether dual-initiation promoter genes are subjected to differential post-transcriptional/translational regulatory mechanisms involving NMD in zebrafish development. To test post-transcriptional regulation of YR and YC initiated RNAs, we blocked translation/NMD in zebrafish embryos by cycloheximide at 22 somites stage for 2 hours until prim 5 stage and performed CAGE analysis (**Figure 6a**). These stages were chosen for the analysis because YC-initiation is broadly active (**Supplementary Figure 1a**; **Figure 3b**) by these stages and maternal mRNAs, which could bias monitoring of post-transcriptional control have been cleared from the embryo^34^. Overall, expression levels of zebrafish *gas5* mildly increased upon cycloheximide treatment with YC-initiation mildly upregulated and YR-initiation downregulated (**Supplementary Figure 6a),** suggesting that *gas5* is regulated by NMD in zebrafish similarly to human yet CAGE-based initiation profile analysis revealed differential response between YR-initiation and YC-initiation. To further demonstrate the response to cycloheximide by individual initiation sites within a single dual promoter, we highlight ribosomal protein (*rps13*) with multiple YR-initiations and YC-initiations (**Figure 6b**). Expression levels of both YC-initiation products are significantly upregulated while YR-initiation products are significantly downregulated (Fisher-exact test; p-value=3.5e-06), suggesting that the intertwined YR-initiation and YC-initiations are independently regulated.

**Figure 6.**
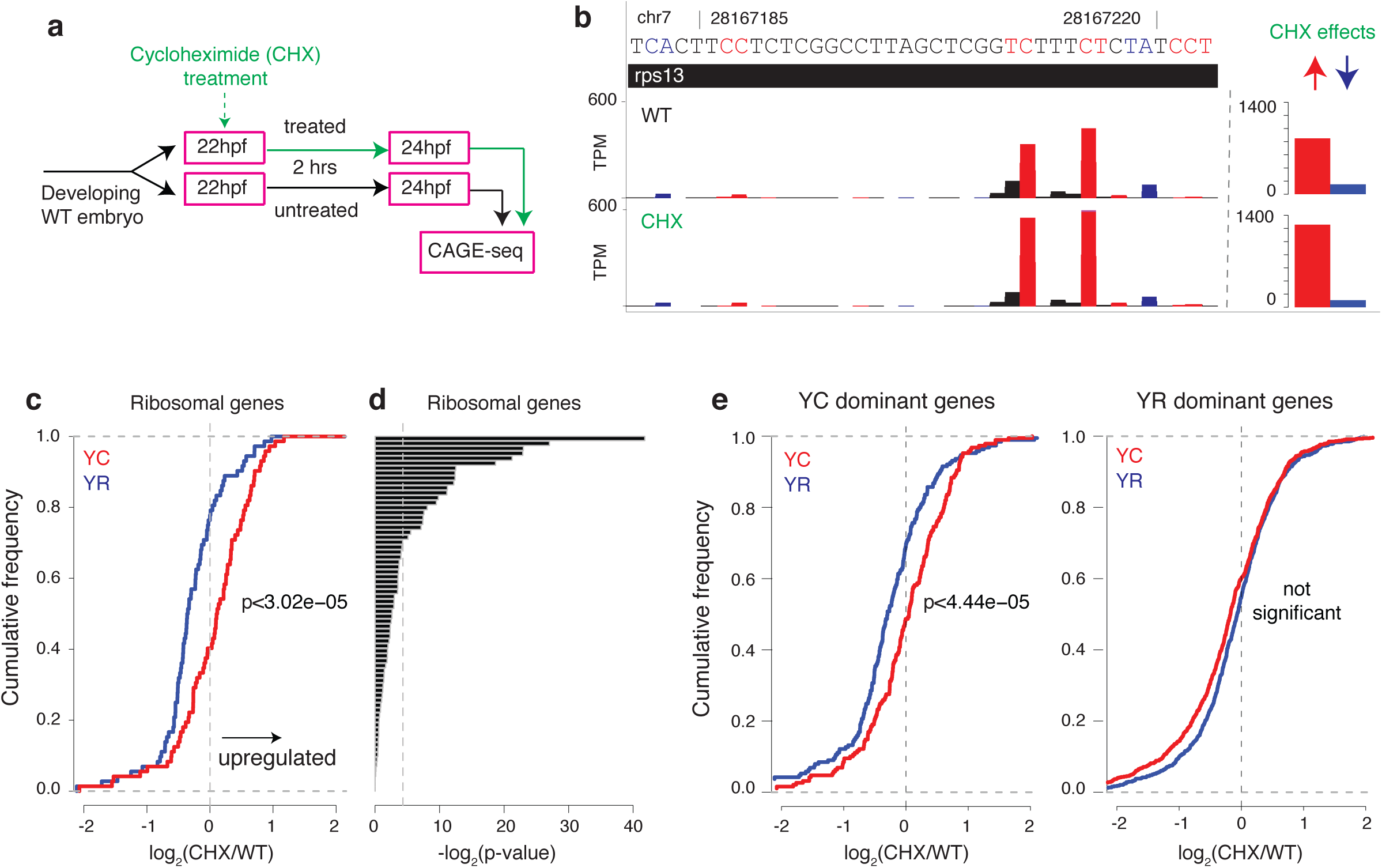
Differential regulation of YR-initiation and YC-initiation during translation inhibition suggest differential translational fates. (**a**) Experimental design to study response of YC-initiation and YR-initiation products during translation inhibition by cycloheximide. (**b**) A UCSC browser screen shot showing an example of levels of YR and YC components of the dual-initiation promoter gene rps13. The bar chart includes sum of all peaks. (**c**) Cumulative frequency of YR-initiation and YC-initiation of all ribosomal protein genes after cycloheximide treatment. X axis indicates the log2 fold change of YR-initiation and YC-initiation in cycloheximide and wild type condition. (**d**) Difference of YR-initiation and YC-initiation in individual ribosomal protein genes after cycloheximide treatment. Each bar represents a ribosomal protein gene. Vertical line represents the significant p-value (0.05) determined by Fisher test. (**e**) Behavior of YR-initiation and YC-initiation in YR-dominant (N=1771) and YC-dominant (N=241) genes.

Next, we expanded the initiation analysis to all ribosomal proteins and subsequently to genome-wide upon cycloheximide treatment. Translation inhibition resulted in an overall upward trend of YC-initiation and downward trend of YR-initiation among ribosomal protein family genes (**Supplementary Figure 6b**). In total, 60% of ribosomal genes have an upregulated YC-initiation while 80% of YR-initiation are downregulated (**Figure 6c; Supplementary Table 5**), corresponding to a significantly different (Kolmogorov-Smirnov test; p=3.5e-06) response between two initiators. However, at the individual gene level only 21 (29.1%) of ribosomal protein genes have significantly different (Fisher’s exact test: p<=0.05) dynamics between YR-initiation and YC-initiation (**Figure 6d, Supplementary Table 5**). Subsequently, we have analyzed the response to cycloheximide for all DI promoter genes by classifying them either as YR-dominant (N=1774) or YC-dominant (N=241) based on the YC/YR expression ratio. YR-initiation and YC-initiation products show significantly different response to cycloheximide (Kolmogorov-Smirnov test; p-value=4.44E-05) in YC-dominant genes (**Figure 6e**) and no significant difference in YR-dominant genes (**Figure 6f**). Taken together, these results demonstrate that the two initiation products differentially regulated the YC-dominant subset of DI promoter genes upon cycloheximide treatment.

### Dual-initiation promoter genes are conserved across metazoans

Finally, we asked whether DI promoters observed in zebrafish are present among other metazoans. We first re-analyzed transcription initiation of the human snoRNA host gene *GAS5* that is transcribed by a 5’-TOP promoter^6^. Visual inspection of combined CTSSs from FANTOM5^22^ revealed that *GAS5* utilizes the expected YC-initiation as dominant initiator (indicated by arrow) (**Figure 7a**) but also an unexpected presence of YR-initiation at a comparable expression level. We measured the expression levels of both initiators in individual cell types across FANTOM5 libraries and observed unexpectedly higher levels of YR component of *GAS5* promoter activity than its YC component in multiple cell types (**Figure 7b**). This result demonstrates the presence and differential expression dynamics of two initiations in a dual-initiation promoter in mammals. We then analyzed DI promoters by adapting the pipeline described in Figure 1a to human HepG2 cell line^22^ and *Drosophila melanogaster* S2 cells^35^. Among expressed genes, 3920 (45%) promoters in HepG2 and 1701 (16%) promoters in S2 cells have intertwined YR-initiation and YC-initiation within the same core promoter (**Figure 7c**). The YC-initiation is dominant in 11.83% and 7.99% of DI promoters in human and *Drosophila* respectively (**Supplementary Table 6**). Furthermore intersection of human and zebrafish orthologous DI promoter genes revealed that 1171 (38.46%) genes share the DI promoter feature indicating high degree of conservation of DI promoters among vertebrates. Gene ontology analysis of DI promoter genes in human has revealed enrichment for translation regulation, mRNA stability, and RNA splicing in human (**Figure 7d**) similar to that in zebrafish (**Figure 2b**) and suggesting that what were previously described as 5’-TOP/TCT promoters, are better described as DI promoters in several cell types both in human and *Drosophila* and argues for redefining non-canonical initiator promoters in these metazoans.

**Figure 7.**
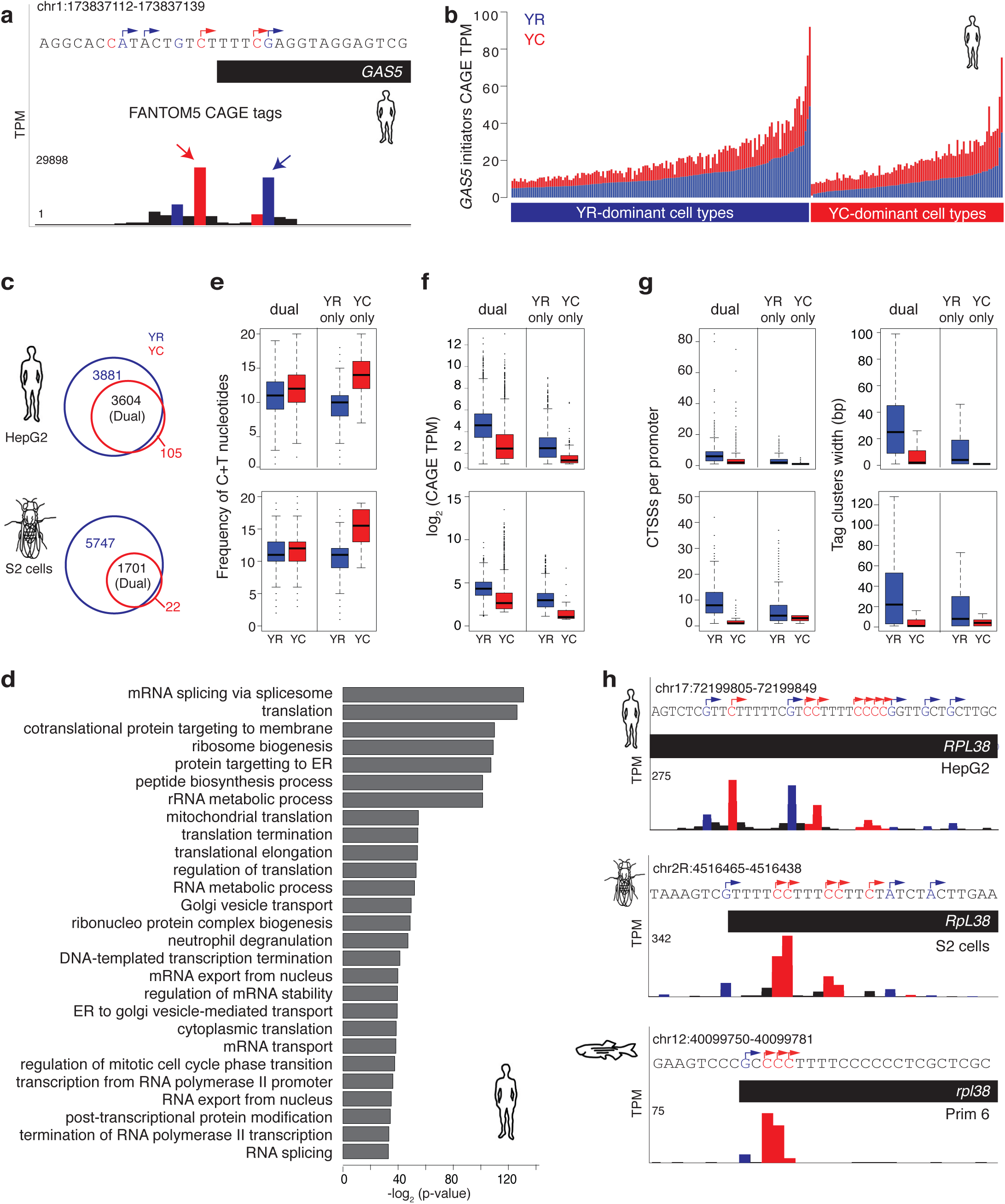
Dual-initiation promoters are conserved in human and Drosophila. (**a**) A UCSC browser screenshot of human GAS5 promoter with FANTOM5 CTSSs summed in hundreds of cell types. CTSSs show transcription of YR-initiation and YC-initiation within same core promoter region. (**b**) Expression levels of YR-initiations and YC-initiations by summing their CTSSs. Promoter are classified as YR-dominant or YC-dominant across individual cell types and their expression is shown in stacked bars. Y-axis shows the expression levels measured in tags per million (TPM). (**c**) Venn diagram with intersection of gene promoters with YR and YC-initiation in human HepG2 and Drosophila S2 cells. Dual-initiation (DI) promoters are indicated in the overlap between detected YR-initiation and YC-initiation. (**d**) Enrichment of gene ontology terms of DI promoters in human HepG2 cell line. (e) Comparison of C+T sequence content around transcription start sites in DI promoters with YR-only or YC-only initiation promoter in human and drosophila. Expression levels of DI promoter genes in human and Drosophila. (**g**) Frequency of CTSSs and promoter width of DI promoters in human and Drosophila. (**h**) UCSC browser screenshots showing CTSSs in the promoter region of RPL38 gene in human, Drosophila and zebrafish. YR-initiation and YC-initiation peaks are colored as blue and red

We next sought to compare sequence content, analyze expression levels and promoter width of dual-initiation promoters in human and *Drosophila*. In both species, DI promoters have higher C+T content around the TSS as compared to YR-only promoters but lower than YC-only promoters (**Figure 7e**), similar to observations in zebrafish (**Figure 2c**). Dual-initiation promoters are highly expressed compared to YR-only and YC-only initiation promoters, which appears to be a shared feature among all three species (**Figure 7f**; **Figure 1d**). Dual-initiation promoters have higher number of CTSSs, resulting in broad promoter shapes, whereas the YC component shows sharp peaks similar to zebrafish (**Figure 7g** compare to **Figure 2g**). The UCSC browser view of the orthologous ribosomal protein genes *RPL38* shows a similar intertwining of YR and YC-initiation events across all three species (**Figure 7h**). Taken together, the above results demonstrate that DI promoters are pervasive and an evolutionary ancient phenomenon characteristic to distant clade with highly conserved promoter architecture and expression features shared among metazoans and highlight the importance of this novel promoter structure organization in divergent animal systems.

## Discussion

In this study, we demonstrate the pervasive nature of non-canonical transcription initiation intertwined with canonical initiation within the core promoter of thousands of genes in zebrafish development. Thus YC-initiation is utilized by a much larger set of genes than previously reported, which was limited to components of translational machinery^6,7,12,17^, and characterized as 5’-TOP/TCT initiators. This dual-initiation arrangement represents a novel composite promoter architecture, which encompasses two sets of targets for transcription initiation in individual promoters. By exploiting the dramatic switch of the embryo transcriptome during the maternal zygotic transition, we show that two initiations are uncoupled from each other during this transition, demonstrating the differential use as well as evidence for lack of interdependence between them in many genes. The apparent independent regulation of initiation site selection in dual promoters during the MZT argues for two initiation mechanisms acting both in the oocyte and the early embryo. However, their use is not selective to ontogenic state, instead it appears to alternate among promoters. The remarkable overlap of transcription initiation mechanisms in the same promoter regions suggest that promoters of dual-initiation genes may respond in more than one ways to regulatory inputs acting in different ontogenic contexts, such as the maternal to zygotic transition (**Figure 8a**).

**Figure 8.**
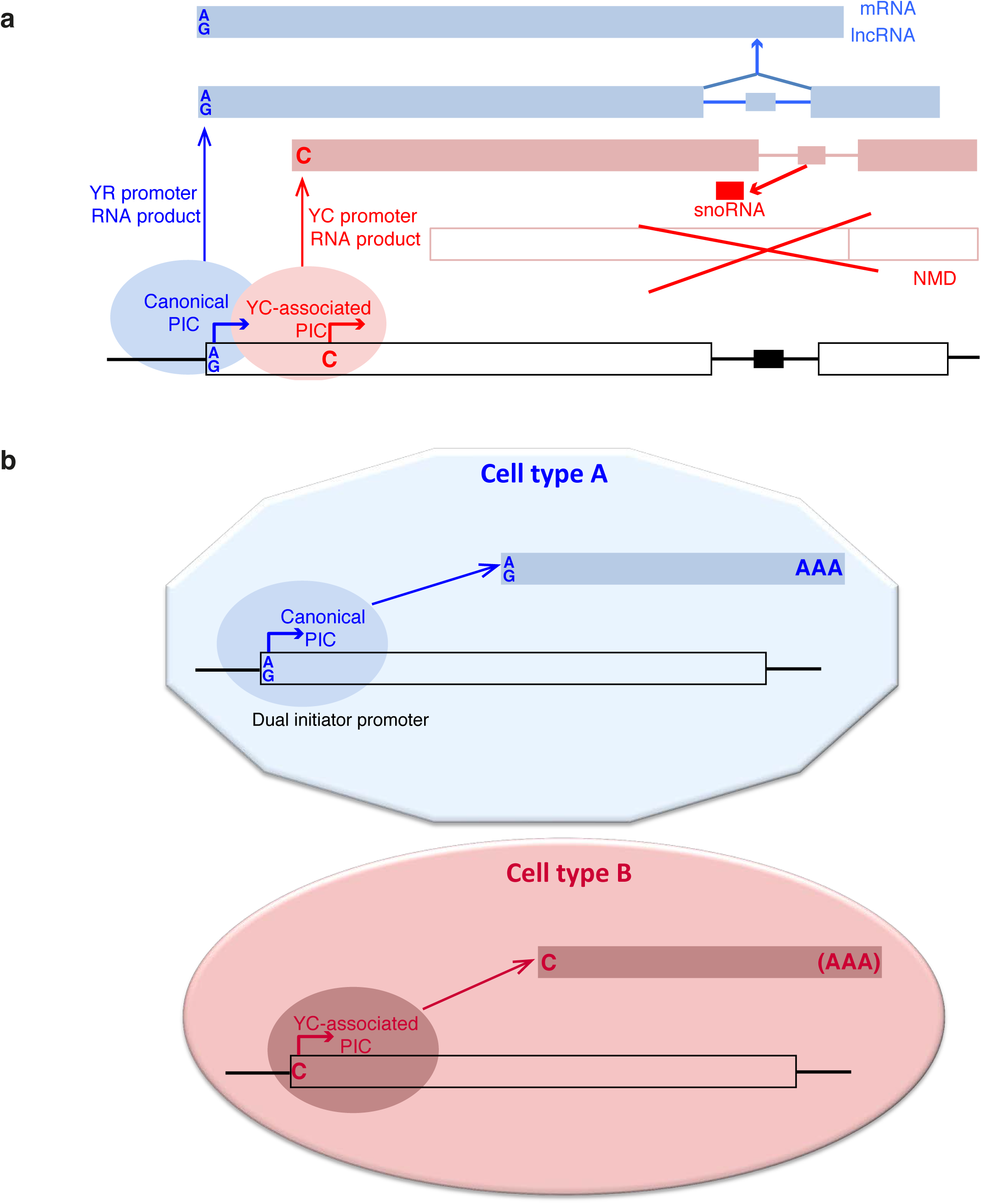
Models for utilization of dual-initiation promoters during development. (**a**) Dual-initiation promoters are occupied by pre-initiation complexes (PIC) in a cell to generate two different RNA products. PIC forms to generate RNA from YR-initiation site for generating a protein coding mRNA or non-coding RNA gene product from a snoRNA host gene, while the YC-initiation may be utilized by a specialized PIC to produce an RNA which is processed to splice out snoRNAs while the rest of the YC initiated RNA subjected to NMD or other degradation pathways. (**b**) Dual-initiation promoter is utilized divergently by YR and YC associated initiation complexes to adapt to requirements in different cells (for example the oocyte versus zygotically active embryonic lineage cells).

We provide evidence that zebrafish snoRNA host genes are transcribed from YC-initiation similar to other model systems^6,7^. However, we also observe that snoRNA host genes also carry canonical YR-initiation not only in zebrafish but also in mammalian cells. While short read sequencing used either in CAGE and RNA-seq is not suitable to directly trace YR- and YC-specific full length RNAs and thus, unequivocally uncouple the post-transcriptionally generated secondary RNA products from two initiation sites. Nevertheless, we show an association of YC-initiation with snoRNA generation by expression correlation analysis of initiation usage. Our results are in agreement with a previous study, which demonstrate that experimentally replacing YC-initiation (5’-TOP) snoRNA promoter with a YR-initiation site reduce snoRNA production^11^. Taken together, our observations strongly argue for a combination of transcription initiation mechanisms acting on snoRNA and host genes and raises the question, whether the mixed nature of canonical and non-canonical initiators reflect a shared promoter region being used by two transcription initiation complexes. Thus a regulatory level at transcription initiation can lead to the production of transcripts with distinct post-transcriptional fates, representing two different functional products, such as snoRNAs and host genes products (see examples of spatial expression of *nanog* derived snoRNA and host gene products in **Figure 5)**. This dual role of a promoter in a single ontogenic stage within the same cell expands the transcript repertoire of the cell (see model in **Figure 8b)**, and if generally applied by dual promoters, could substantially impact on the a yet unexplored additional layer of diversity of RNAs produced from genes. We hypothesize, that the expansion of utilization of a non-canonical initiation to a wide range of genes could indicate a general transcription regulation paradigm, which represents adaptation to differential regulation of a variety of promoters^15,18^. Dual-initiation promoter genes are highly expressed compared to other genes (**Figure 1d; Figure 7f**), which is not specific to the contributing YC components, as expression levels of the corresponding YR component alone is also higher than that of YR-only or YC-only initiator genes. This observation either suggest that sharing two alternative initiation mechanisms leads to boost of expression levels or suggest that YC-initiation might be evolutionary co-opted in highly expressed genes. It is interesting to note that the efficiency of transcription correlates positively with translation efficiency and raises the possibility that highly expressed DI promoters contribute to coordination between transcription and translation^36^. The enrichment of translation and RNA regulation related gene ontology terms in DI promoter genes, along with notable absence of developmental regulator genes, raises the question of why and how this promoter architecture evolved. Important insight into potential functional significance of the non-canonical initiation comes from studies on target genes of the mTOR pathway that are translationally regulated^15,16^, and are enriched in 5’-TOP/TCT initiator. Polypyrimidine proximal to 5’ end of these genes is a target for translation regulation and has been suggested to serve as a targeting mechanism for oxidative and metabolic stress or cancer induced differential translation regulation by the mTOR pathway^15,16,18-20,37^. Other studies argue for the co-transcriptional regulation of post-transcriptional fates of RNAs, where promoter identity influences cellular localization and translation efficiency of mRNAs under different environmental conditions^38,39^. Thus, it is plausible that specialization of transcription initiation has co-evolved with post-transcriptional regulation to regulate RNA fates by transcription and the 5’-ends of TOP RNAs reflects such a dual regulatory function. Dual-initiation promoters offer the potential for linking translational regulation to transcriptional regulation in a large range of genes and thus increase the repertoire of genes that may respond to such signals. In this study we have identified many genes, which carry low level of YC-initiation events, which may reflect a non-induced ground state for YC regulation. However there was a notable correlation between the length of polypyrimidine stretch at the 5’ end and the expression level of YC (**Figure 2e**). It is not yet possible to distinguish in the CAGE dataset whether this correlation reflects RNA stability or transcriptional differences. Nevertheless, an unanswered question remains, whether the polypyrimidine stretch at the 5’-end is required for selective translation factor binding such as eIF4F complex or also represent distinct transcription regulatory signals acting at the transcription initiation level.

The current definition of 5’-TOP mRNA includes a stretch of minimally 4 to 13 pyrimidine^17^ based on observations restricted to translational-associated genes^17^, which have longer pyrimidine stretch also in zebrafish (**Supplementary Figure 2d**). This definition has been suggested to be potentially too stringent, as translationally regulated genes revealed by ribosome profiling are enriched in transcription initiation with “C” and carry only a short pyrimidine stretch^15,16^. We used a threshold of 1 TPM and identified thousands of YC-initiation sites and thus expanded the pool of genes, which ought to be considered when transcriptomic responses to metabolic stress for example via the mTOR pathway are sought and our results argue for the need for discriminating RNAs produced from the same promoter by using transcriptome analyses with single nucleotide resolution. Many of these genes may respond to post transcriptional signals similarly to 5’-TOP promoter genes, however this response is potentially masked in investigations in which the RNAs with distinct initiation profiles are not separately quantified. This possibility is demonstrated by our cycloheximide treatment experiments where YR and YC components of dual-initiation promoters respond differentially to interference with translation/NMD, which implies that C-capped transcripts may be more prone to NMD than their A/G-capped counterparts originating from the same DI promoter. Taken together, our findings provide a framework for future studies to understand coordinated regulation of transcription and translation of thousands of genes.

The unexpected widespread presence of YR and YC-initiation intertwined in the same core promoter raises a question as to why this pervasiveness was not seen before. Previous studies analyzing TSSs in a genome-wide level reported multiple TSSs in same core promoter^2,3,5,22,26^, but downstream analysis is focused on dominant TSSs, majority of which appear as YR, and as a result YC-initiation remained unexplored. Reinvestigation of human and *Drosophila* cell line datasets in this study demonstrated that the dual-initiation is a widespread phenomenon and share similar sequence feature, promoter shapes, expression levels and enriched gene ontology. Dual-initiation promoter genes in three major metazoan model systems spanning a very large evolutionary distance and across many orthologues suggest an evolutionary ancient shared promoter architecture with fundamental functions in multicellular function and development and motivates future investigation into the functional consequences of selective transcription initiation within gene promoters in general. Our studies in zebrafish embryos with dynamic spatiotemporal transcriptional patterns underscore the importance of further analysis of the dynamics of YR and YC expression profiles across multiple cell types and varying physiological states in other model systems.

## Materials and Methods

### Zebrafish CAGE data after cycloheximide treatment

We generated zebrafish CAGE data for translation inhibition experiment. Zebrafish embryos were treated with 100 µg/ml cycloheximide (Sigma-Aldrich) or 0.1 % DMSO as control for 2 hours, starting at 22 hours post fertilization (hpf). Total RNA was extracted from the control and treatment groups at 24 hpf using TRIzol (Invitrogen/ThermoFisher) following the manufacturer’s instructions and used for CAGE libraries preparation as described before^3^, except for the use of oligo-dT primer instead of random primers in the first strand synthesis step. CAGE libraries were sequenced on Illumina MiSeq system.

### RNA sequencing of capped RNAs

Total RNA was extracted from 24 hpf embryos using TRIzol reagent (ThermoFisher) and DNAse treated using TURBO DNA-free™ Kit (ThermoFisher) according to the manufacturer’s instructions. Full length cDNA libraries were prepared using TeloPrime Full-Length cDNA Amplification Kit (LexoGen), designed to capture 5’ Capped, polyadenylated transcripts. Two full cDNA libraries were prepared (technical replicates) according to the provided user manual, using 2 µg of total RNA as input, with differing numbers of PCR amplification cycles: 14 and 16 respectively. Sequencing libraries were prepared from both cDNA libraries using the MicroPlex-Library-Prep-Kit-v2 (Diagenode) and sequenced (2×100bp reads) on HiSeq 2500 System (Illumina). For identification of transcription start sites, only reads starting with the 5’ TeloPrime adapter were selected, trimmed using cutadapt^40^ and mapped to the zebrafish Zv9 zebrafish reference genome and Ensembl version 79 transcript annotations using STAR^41^, reporting only uniquely mapped reads. CAGE-like TSS (ClTSS) were called using CAGEr package^42^. ClTSS with at least 0.3 tpm were assigned to Ensembl version 79 promoter regions (500 bp upstream and 250 bp downstream from the annotated transcript start). A given promoter was identified as YR or/and YC-initiation type, if the mean sum (of the two technical replicates) for the corresponding ClTSS-initiator signals (YR/YC) was at least 3 tpm, i.e. the same criteria used for CAGE samples.

### Publicly available CAGE data on zebrafish, human and fruit fly

CAGE data on zebrafish, human and drosophila were downloaded from previous studies. Mapped zebrafish CAGE data was used from previous study^3^. Mapped human CAGE data was downloaded from FANTOM5^22^. Three replicates of HepG2 CAGE data was merged and converted CAGE tags count into tags per million (TPM). Drosophila CAGE raw reads was downloaded from modENCODE^35^. CAGE libraries were mapped using bowtie2^43^. We allowed two mismatches and only unique mapping reads were retained. Mapped reads having a “G” mismatch in the first nucleotide was corrected and transcription start site was corrected accordingly.

### Downstream analysis of CAGE data

Based on −1 and +1 nucleotides for each CAGE Transcription Start Site (CTSS) we classified Y_-1_R_+1_ (Y: pyrimidine (C/T)) and (R: Purine (A/G)) as canonical initiator^2,3^ and Y_-1_C_+1_ as non-canonical initiator. For all analysis, we selected CTSS with a minimum expression level of 1 tag per million (TPM) in one of the 12 developmental stages. From the above pool of selected CTSSs, we intersected *remaining* CTSSs and included those CTSS with a minimum of 0.5 TPM. Canonical and non-canonical initiators were separately clustered if they overlapped within 20 nucleotides in the same strand resulting a tag clusters (TCs). Expression levels of all CTSS falling within the tag clusters are summed that gives the expression level of tag clusters. CTSS with the highest expression level, within the tag cluster, defines the dominantly used transcription start sites. The width of tag clusters defines promoter shape which is classified as sharp or board. Genes expression levels are calculated by aggregating tag clusters in the assigned promoter region (500 nucleotides upstream and 300 nucleotides downstream of Ensembl annotated TSSs). Canonical and non-canonical expression levels of each gene were calculated by separately aggregating canonical and non-canonical CTSS.

### Annotation of zebrafish snoRNAs from size selected small RNA reads

Size selected (18-350 nucleotide) zebrafish small RNA-seq data was downloaded from public dataset^23^. Adapters were filtered, and mapped sequence reads to zebrafish genome (zv9) using bowtie2^43^. Sequence reads were first mapped to ribosomal RNAs (rRNAs) and excluded those mapping to rRNAs. Unmapped reads were then remapped to genome by allowing up to four multi mappings reads. To ensure that snoRNAs are annotated from mapped reads that resemble the expected full-length of snoRNAs, we retained only those mapped reads that longer than 50 nucleotides and potentially represent full-length snoRNAs rather than small RNA fragments. SnoRNAs were annotated by using four different tools, namely Infernal^44^, snoReport^45^, snoGPS^46^ and snoscan^47^. Infernal was used together with covariance model from RFAM^48^. An e-value cutoff of 0.05 for each covariate model provided by RFAM was used. SnoReport, snoscan and snoGPS were used with default parameters for annotation of novel snoRNAs. To retain high confidence snoRNAs, we excluded snoRNAs that have low reads (<5 reads), residing on exons and repeats. Ensembl (version-79) has 312 annotated snoRNAs^49^ and 270 of them are supported by at least 5 reads in developmental stages we analyzed. Out of 270 snoRNAs from Ensembl, we predicted 264 snoRNAs and annotated 176 novel snoRNAs. We finally quantified snoRNAs expression by counting mapped reads using BEDTools^50^. Total mapped reads were calculated using SAMtools^51^ and then converted into reads per million.

### Gene Ontology

Gene Ontology analysis was done by using GOstats package^52^ from BioConductor^53^. Over-represented GO terms were corrected for multiple testing with the Benjamini-Hochberg false discovery rate and obtained statistically significant GO terms by applying a p-value cutoff of <= 0.05.

### Data visualization

A genome browser view of multiple genes was downloaded from UCSC genome^54^ CTSSs and other relevant data were uploaded on UCSC Genome Browser as tracks for visualization. A screenshot of promoter regions with data tracks were downloaded from the UCSC browser. All other figures were made using R.

### RNA extraction and RT-PCR amplification

Purification of total RNA was performed using miRNeasy mini kit (Qiagen, Cat. 217004) following the manufacturer instructions. cDNA was synthesized using the iScript cDNA synthesis kit (BioRad, Cat. 170-8890) from 200ng of purified RNA and snoRNA sequences were amplified by RT-PCR. Amplified cDNAs were verified by electrophoresis in 4% MetaPhor agarose gel (Lonza, Cat. 50184). We used the following primers for amplification: ***dkc1*-snorna**: TGATGAACTTGTTTATCCATTCGC and TGTCAGTCATGTATAATCATCTTGGC; ***nanog*-snorna**: CGTGTCCATGCTGTTGCTTG and CTTGTATCATCGTGCCTTTAAGACG.

### Riboprobes, single and double fluorescent whole-mount in situ hybridization

T3 promoter was linked at the 5’ and the 3’end of the full-length cDNA for each amplified snoRNAs for the synthesis of antisense and Sense riboprobes, respectively. Transcription were done by T3 polymerase using digoxigenin (DIG) labelling mix (Roche) or DNP-11-UTP (TSA™ Plus system, Perkin Elmer) according manufacturer’s instructions. The probes were subsequently purified on NucAway spin columns (Ambion), and then ethanol-precipitated. Single whole-mount *in situ* hybridizations were performed as described previously^55^. Double fluorescent *in-situ* hybridizations were carried out as described previously^56^.

### Whole mount immunofluorescence after ISH hybridization

Embryos were washed in wash buffer (PBS, 0.3% v/v triton), incubated in blocking buffer (PBS 1x, Tween 0.1%, Goat serum 4%, BSA 1%, DMSO 1%) for 3 hours and then incubated with primary antibody over night at 4C (Anti-Fibrillarin, Abcam 38F3, 1:10). Embryos were then washed in wash buffer and blocked 3 hours followed by incubation with the secondary antibody overnight at 4C (Anti-Mouse Alexa 633, 1:500).

### Imaging

Microscopy images were obtained with an Olympus DP70 camera fixed on a BX60 Olympus microscope. Confocal imaging was performed using a Leica TCS SP5 inverted confocal laser microscope (Leica Microsystems, Germany) Digitized images were acquired using a 63X glycerol-immersion objective at 1024X 1024 pixel resolution. Series of optical sections were carried out to analyse the spatial distribution of fluorescence, and for each embryo, they were recorded with a *Z*-step ranging between 1 and 2 μm. Image processing, including background subtraction, was performed with Leica software (version 2.5). Captured images were exported as TIFF and further processed using Adobe Photoshop and Illustrator CS2 for figure mounting.

## Data availability

CAGE and RNA-seq data tracks can be visualized and downloaded from the DANIO-CODE DCC (https://danio-code.zfin.org) and the UCSC genome browser public track hub (Promoterome).

## Authors contributions

C.N. and F.M. conceived and coordinated the project. C.N. and Y.H. analyzed data. C.N. and F.M. interpreted results with critical comments from B.L., and J.B.A. E.T.S. R.C. and B.P. designed and performed whole mount in situ hybridization experiments, for which E.T., F.M and B.P. interpreted results. P.C. and A-M. S generated CAGE libraries from cycloheximide-treated embryos. Y.H. performed cycloheximide and RNA seq experiments. C.N. and F.M. wrote the manuscript with contribution from B.L. and J.B.A. All authors read and approved the manuscript.

## Acknowledgements

The authors are grateful to R. Taylor Raborn, Robin Andersson, Pawel Grzechnik, Laszlo Tora and Christian Kroun Damgaard for critical comments on manuscript. This work was supported by the BBSRC (BB/L010488/1) and a Wellcome Trust Investigator award to FM and BL. The laboratory of JBA is supported by the Novo Nordisk Foundation (14040) and Danish Medical Research Council (4183-00118A).

---

**Supplementary Figure 1.**
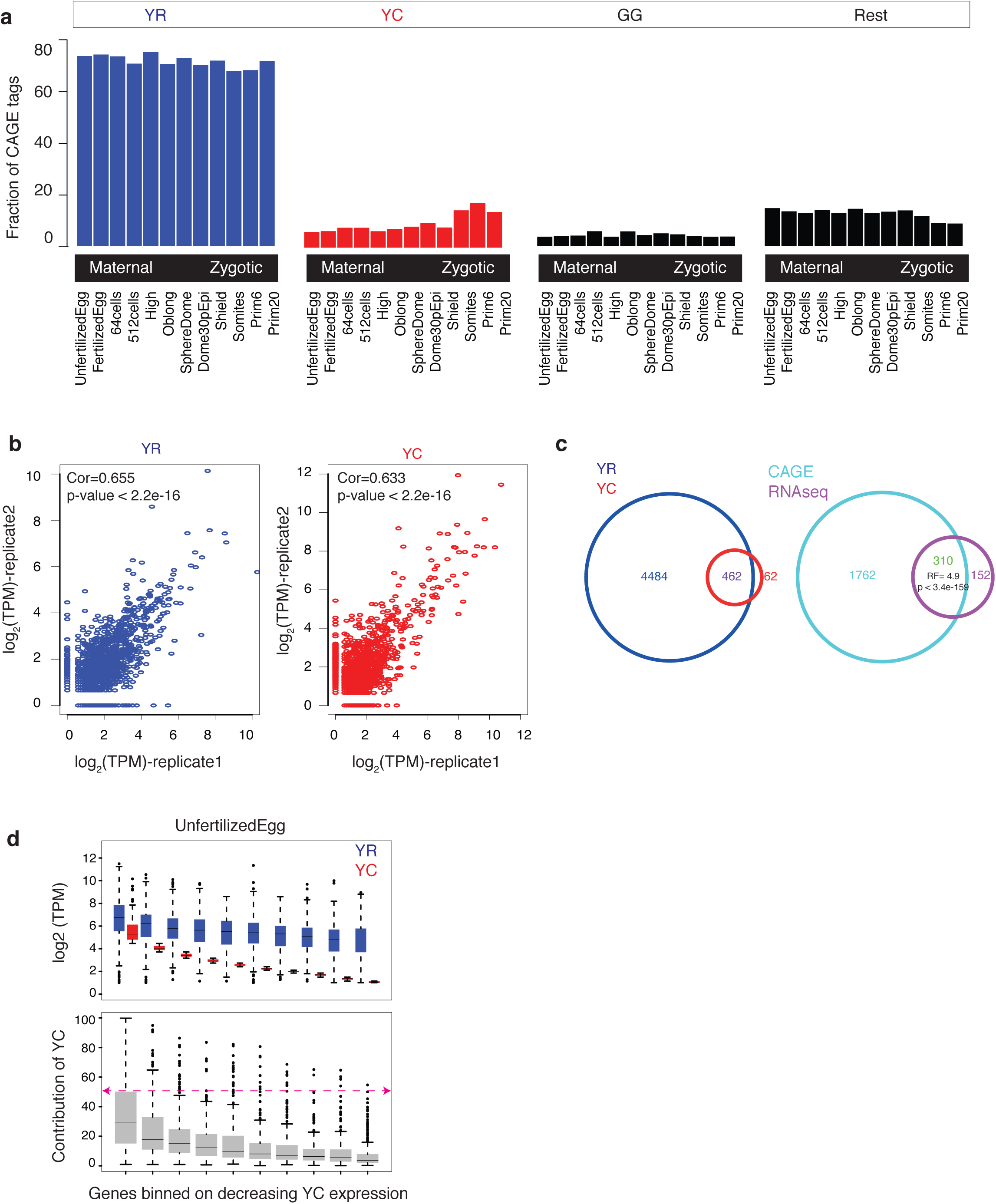
Distribution and correlation of YC-initiaions and YR-initiations in zebrafish developmental transcriptomes. (**a**) Classification of CTSSs based on dinucleotide frequencies around CTSSs Y-axis indicates fraction of CTSSs. (**b**) Correlation of canonical YR-initiation and non-canonical YC-initiation between two replicates of prim 5 stage. (**c**) Intersection of genes with YR-initiation and YC-initiation from 5’ end capped RNA-seq (left). Intersection of dual-initiation genes from RNA-seq and CAGE-seq (right). (**d**) Contribution of YC-initiation with respect to YR-initiation expression levels in unfertilized egg stage. Genes are sorted according to YC expression levels and grouped into 10% bins.

**Supplementary Figure 2.**
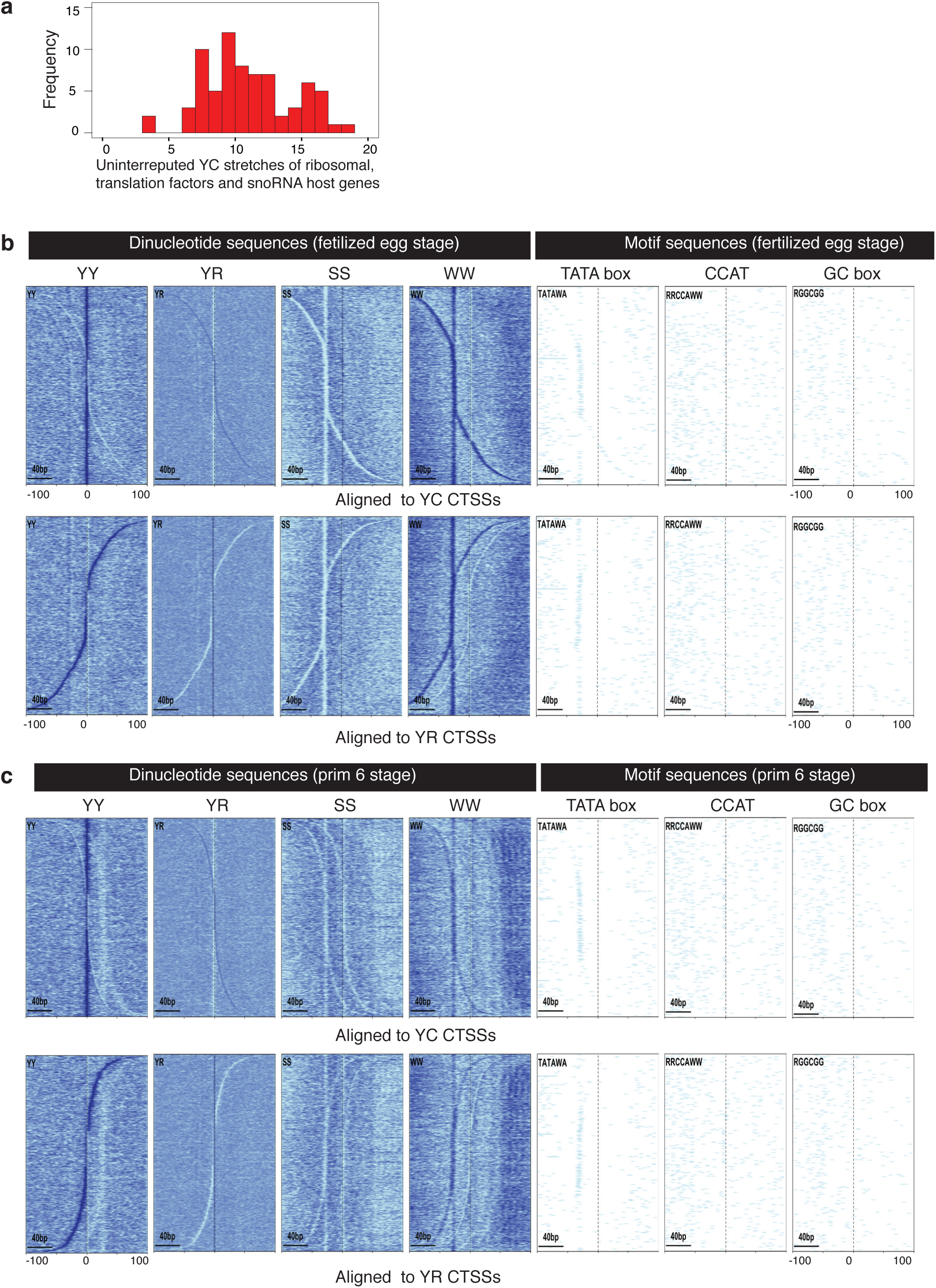
Features of dual-initiation promoter genes. **(a)** Frequency of uninterrupted polypyrimidine stretches around YC-initiation sites of translational-associated genes (ribosomal proteins, translation initiation/elongation factors and snoRNA host genes). X axis indicate the maximum length of uninterrupted stretches of pyrimidine sequence. (**b-c**) Distribution of dinucleotide (YY/YR/SS/WW; Y=C/T; R=A/G; S=C/G; W=A/T) sequence content and (TATA, CCAT and GC box) motifs with respect to YR-initiation and YC-initiation of dual-initiation promoters in (b) fertilized egg and (c) prim 5 stage. Genes are aligned based on distance between YR and YC and aggregated to the +1 position of YR and YC dominant CTSS respectively.

**Supplementary Figure 3.**
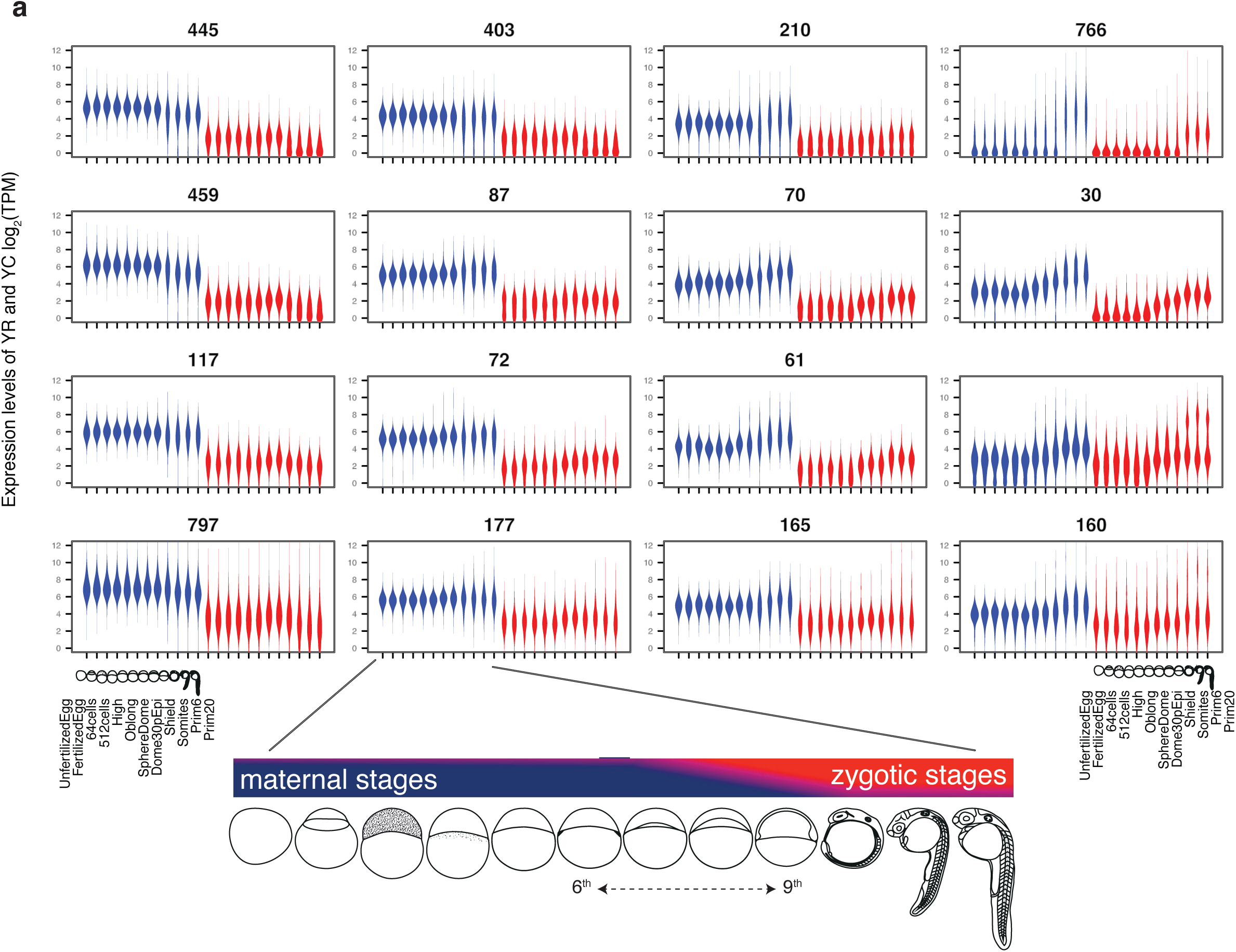
Expression dynamics of YR-initiation and YC-initiation during zebrafish embryo development. **(a)** Self organizing map clusters of the TPM expression profiles of YR and YC components of genes during maternal and zygotic stages. Developmental stages along the x-axis are shown at the bottom. Y-axis indicates the expression levels. Blue and red colors indicate YR and YC components, respectively. Numbers above panels indicate the number of genes in each cluster.

**Supplementary Figure 4.**
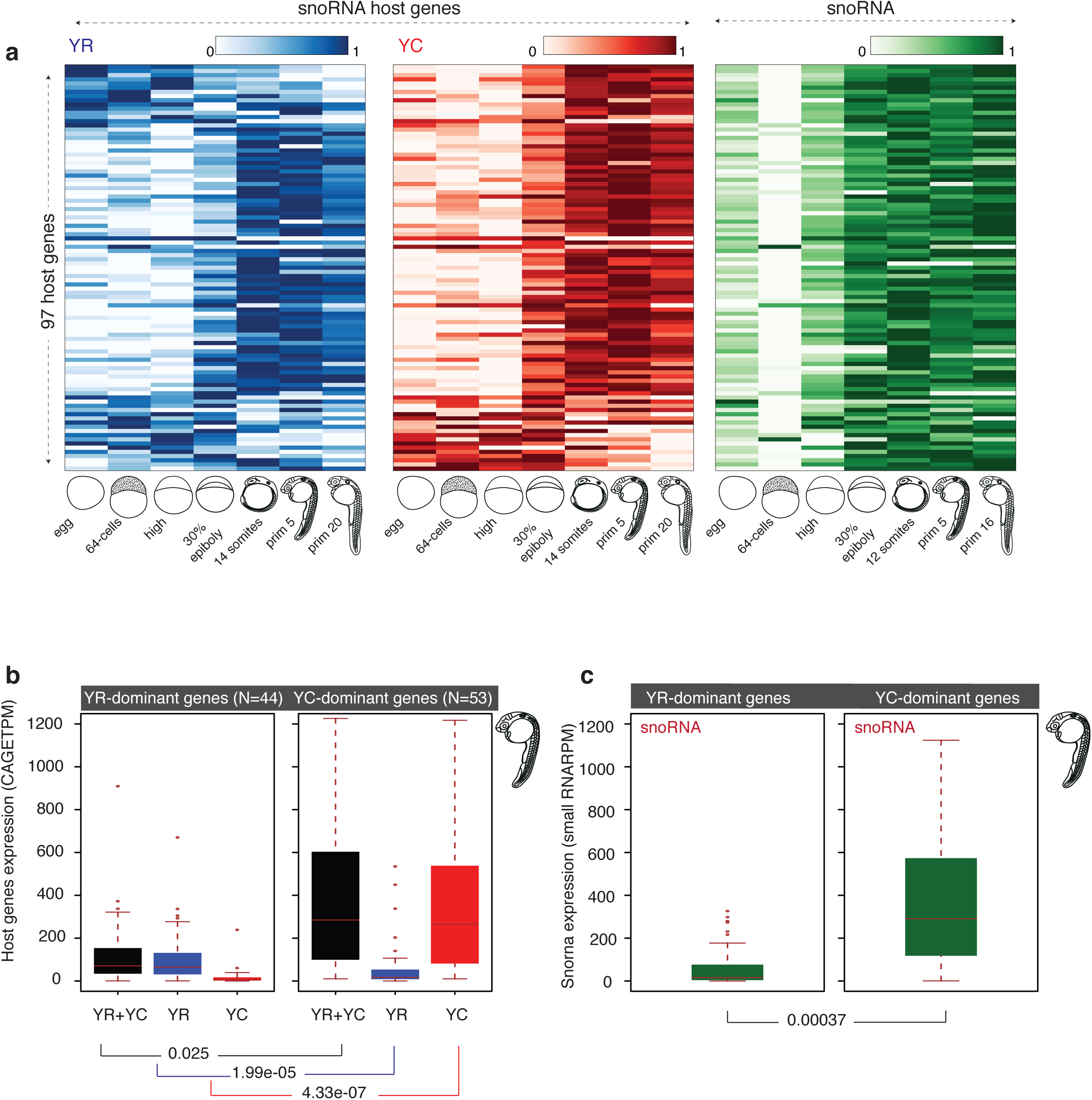
Correlation of YR and YC components of snoRNA host genes with snoRNA expression levels. **(a)** Heat maps showing the expression levels of YR and YC components of snoRNA host genes along with snoRNA expression levels. Genes are sorted based spearman’s correlation values between YR-initiation and YC-initiation. Expression levels are scaled between 0-1 for each row separately for YR, YC and snoRNA. (**b**) Expression levels of YR (blue) and YC (red) initiators, along with combined (black) expression levels of YR and YC. Host genes are divided into two groups (YR-dominant or YC-dominant) based on dominant expression of initiators. Y-axis indicate tags per million. (**c**) Expression levels of snoRNAs transcribed from YR and YC-dominant genes. Y-axis indicate reads per million.

**Supplementary Figure 5.**
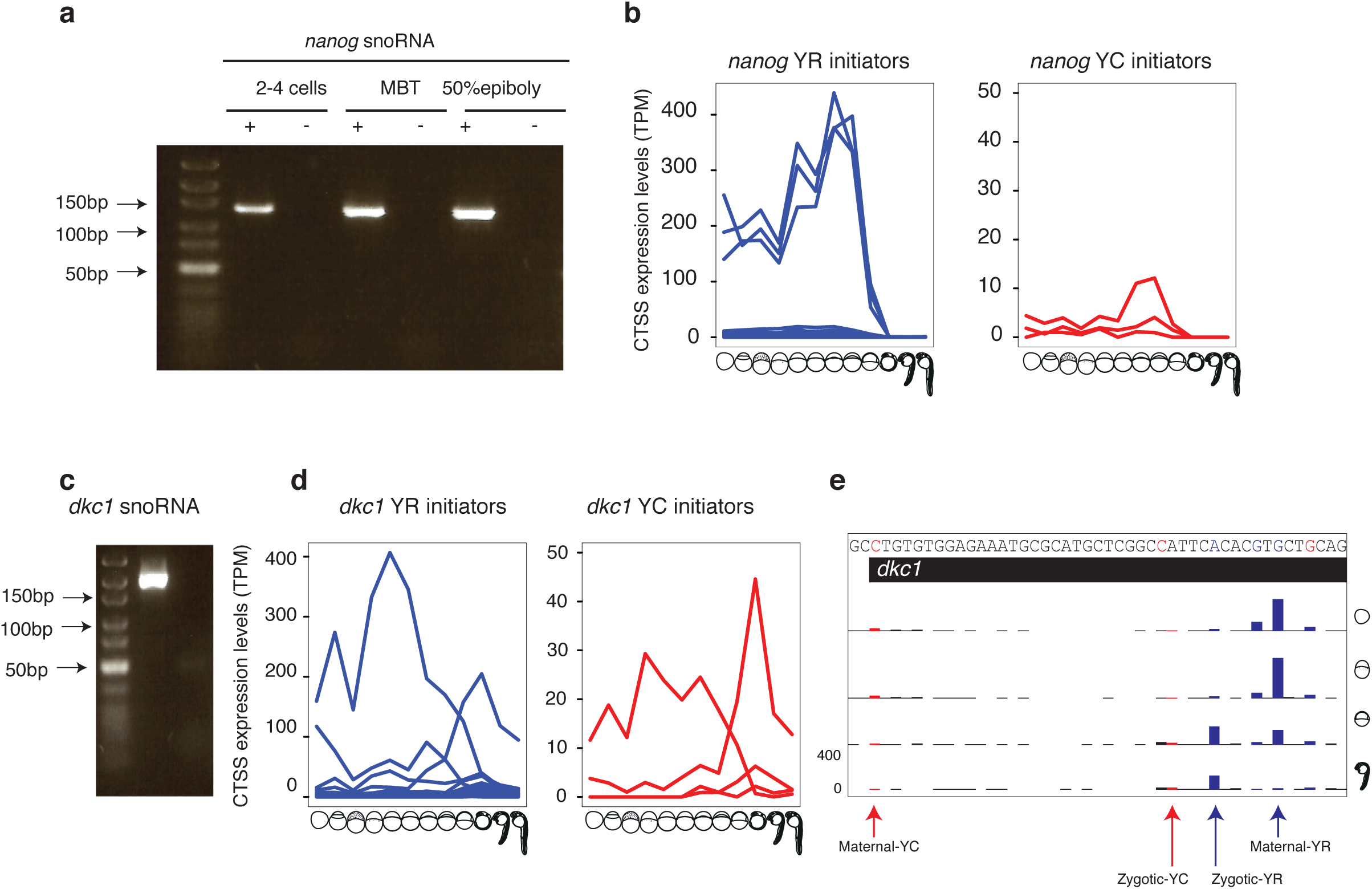
Quantitation and dynamics of snoRNAs and YC-initiations and YR-initiations of their host genes. **(a)** Validation of nanog snoRNA expression by RT-PCR in three developmental stages. Predicted size of PCR fragment is 131 bp (**b**) Expression level and developmental dynamics of individual YR-initiation and YC-initiation in the nanog promoter region. X-axis indicate the developmental stages. (**c**) Validation of dkc1 snoRNA expression by RT-PCR in prim 5 stage. (**d**) Expression level and developmental dynamics of individual YR-initiation and YC-initiation in the dkc1 promoter region. X-axis indicate the developmental stages. (**e**) A UCSC browser screen shot of dkc1 gene with CTSSs. YR-initiation and YC-initiation are colored blue and red respectively.

**Supplementary Figure 6.**
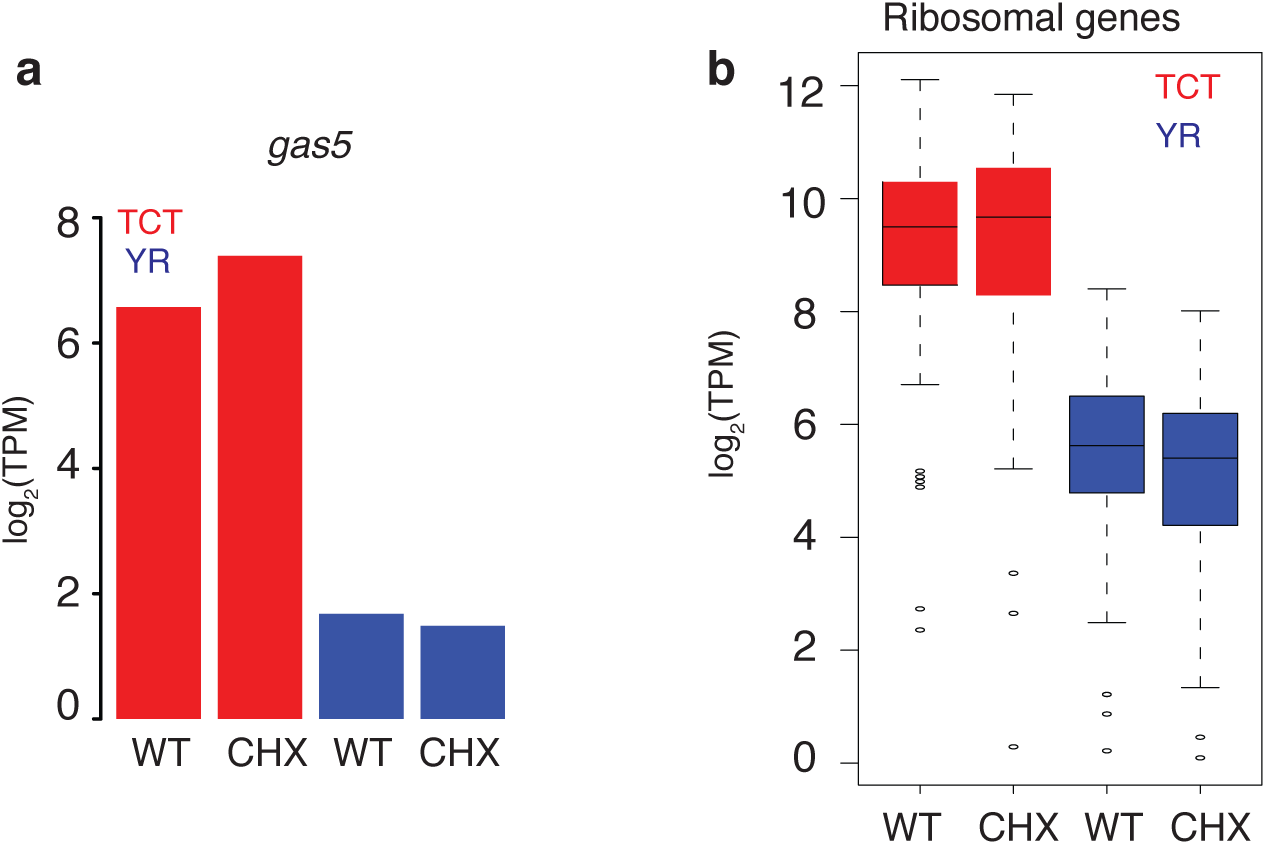
Effect of translation inhibition on YR-initiation and YC-initiation products of dual-initiation promoters. (**a**) Expression levels of YR-initiation and YC-initiation products of gas5 after cycloheximide treatment. Blue and red color indicates YR-initiation and YC-initiation respectively. (**b**) Expression dynamics of YR-initiation and YC-initiation of all ribosomal protein genes.

